# Molecular Determinants Governing the Antitubercular Activity of Griselimycin

**DOI:** 10.64898/2026.03.19.712639

**Authors:** Andrew Spira, Sydney M. Williams, Rachita Dash, Sarah E. Newkirk, Irene Lepori, Yuanhang Caroline Luo, Vaskar Chowdhury, Sobika Bhandari, Zichen Liu, Jean Chmielewski, M. Sloan Siegrist, Marcos M. Pires

## Abstract

The emergence of drug-resistant strains of *Mycobacterium tuberculosis* makes anti-tubercular drug development a critical priority. Griselimycin is a cyclic peptide that targets the essential DNA sliding clamp of *M. tuberculosis*. While griselimycin is a promising antimycobacterial compound, its poorly understood structure-activity relationship has stalled derivatization. To investigate the contribution of each amino acid towards its activity, we assessed the antibiotic activity of an alanine scan library in *M. tuberculosis*. Incorporation of an *N*-terminal fluorophore enabled visualization of griselimycin accumulation inside of mycobacteria-infected macrophage cells. Derivatives containing altered cyclization chemistries, unnatural amino acids, expanded proline side chains, and alkyne handles were assessed, with a final design combining the most potent edits showing increased antibiotic activity. These findings present the comprehensive structure-activity investigation of griselimycin and can be used to rationally design future analogues.

**TABLE OF CONTENTS GRAPHIC:** 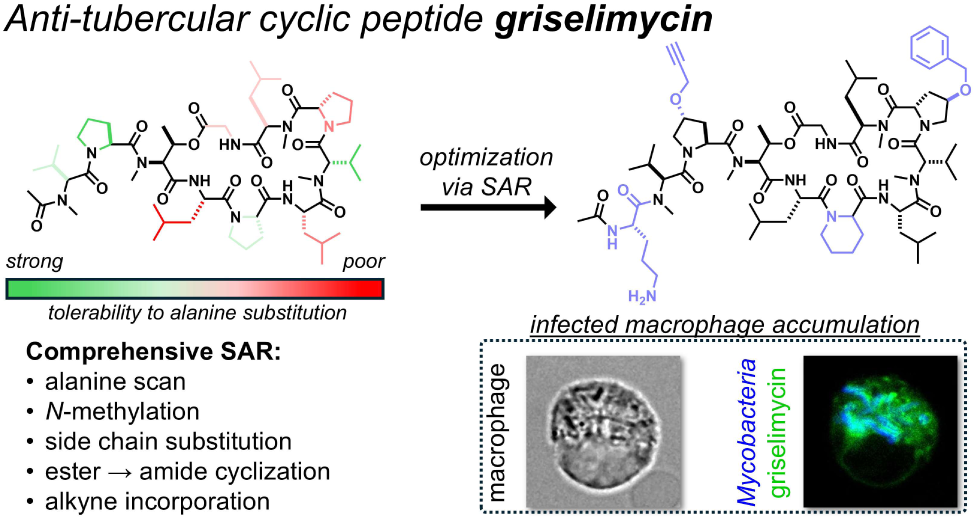

## INTRODUCTION

Tuberculosis (TB) is one of the world’s deadliest infectious diseases, with over ten million cases and one million deaths reported in 2024.^1^ Multidrug-resistant (MDR) strains of the causative agent *Mycobacterium tuberculosis*, while associated with only ∼4% of annual infections, account for nearly 15% of all TB-related mortalities.^2^ The rise of MDR-*M. tuberculosis* and FDA approval of only three new anti-TB drugs in the last 50 years^3^ has led the World Health Organization to designate it as a critical priority for antibiotic development.^4^ Understanding the key structural elements that govern anti-tubercular activity is essential for the rational design of next-generation therapeutics capable of overcoming existing resistance mechanisms.

Griselimycin (**GMe**), first isolated from *Streptomyces griseus* in the 1960s, is a ten-residue cyclic peptide and potential antibiotic that targets the DNA sliding clamp (DnaN) of *M. tuberculosis* (**Fig. 1a**).^5^ **GMe** consists of an eight-residue ring, cyclized via an ester bond between the side chain of Thr_3_ and the *C*-terminus of Gly_10_, and a two-residue tail with an acetylated *N*-terminus. **GMe** contains four *N*-methylated resides (Val_1_, Thr_3_, Val_7_ and D-Leu_9_) and two 4-methyl-prolines in positions 2 and 5, and is thought to inhibit DNA replication by noncovalently binding to the DnaN subunit of *M. tuberculosis* pol III (**Fig. 1b**).^6,7^ DnaN (also called the β subunit) has been identified as a promising antibiotic target due to its essential biological role in DNA replication and poor sequence homology relative to the eukaryotic DNA clamp.^8,9^ DnaN is a torus-shaped homodimer that encircles DNA and complexes with the α (nucleotide-adding) and ε (proofreading) subunits of DNA polymerase III (pol III) during DNA replication (**Fig. 1c**).^10^ Structural alignment of **GMe** co-crystallized DnaN with a pol III holoenzyme suggests that the peptide inhibits assembly of the polymerase complex by occupying the α and ε binding pockets (**Fig. 1d – e**), shutting down DNA replication.^11,12^ Although initial *in vitro* results were promising, **GMe** exhibited poor pharmacokinetics in human trials and was soon eclipsed by other TB drug candidates.^13^ The current first-line treatment regimen for TB includes rifampicin and isoniazid, both of which are ineffective against MDR-*M. tuberculosis* strains.^14–16^ Consequently, **GMe** has seen renewed interest as a failed *M. tuberculosis* antibiotic with potential for improvement.

**Figure 1.**
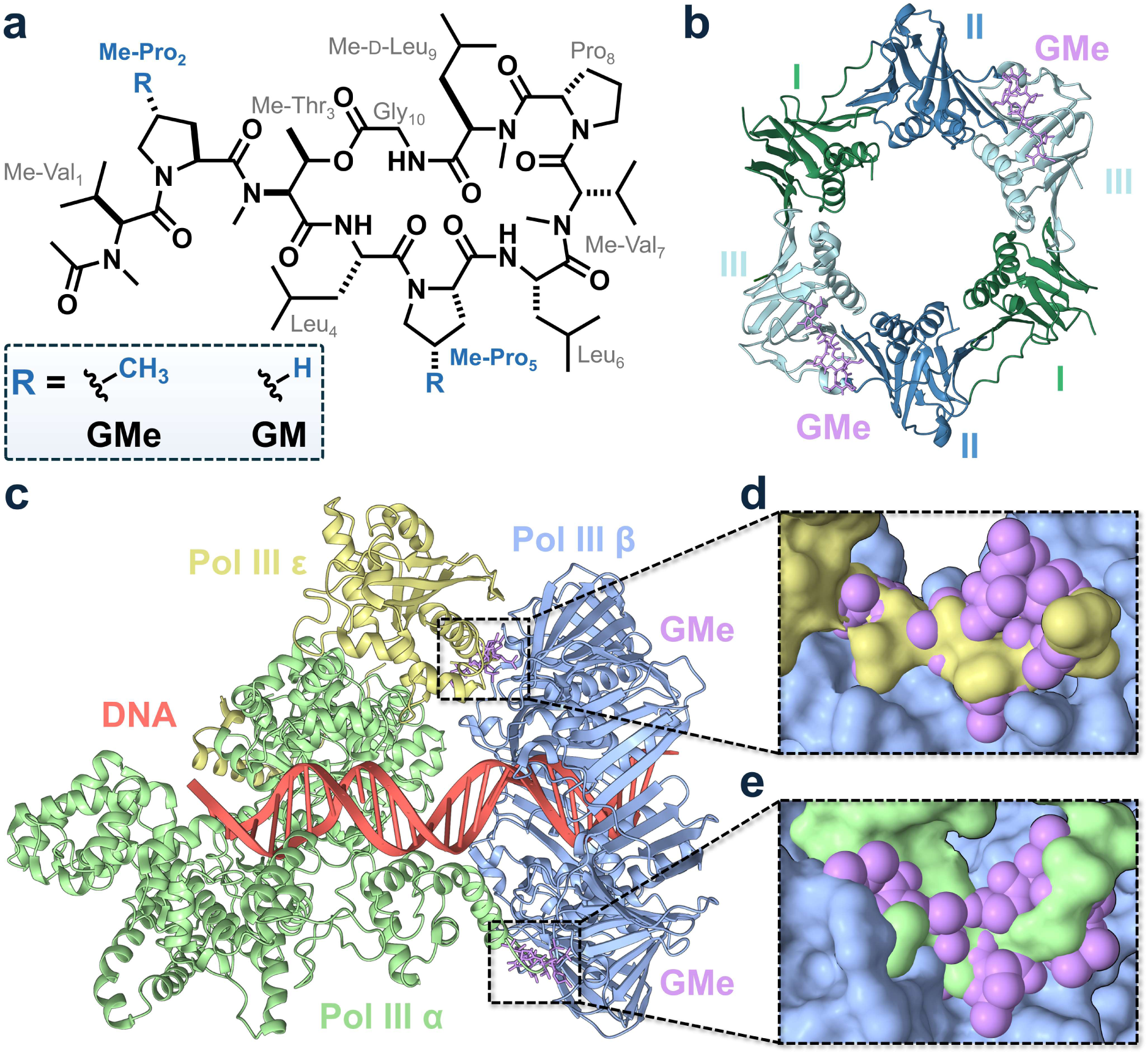
**(a)** Structure of the natural product griselimycin (**GMe**) and a derivative replacing 4-methyl-Pro_2_ and 4-methyl-Pro_5_ with standard proline (**GM**). **(b)** Structure of *M. tuberculosis* DnaN dimer co-crystallized with **GM** (PDB: 5AGU). Domains I, II, and III of each subunit are colored green, blue, and teal respectively. **(c)** DNA Pol III complex from *E. coli* (PDB: 5FKW)^11^ consisting of DNA (red), α (green), β (blue), and ε (yellow) subunits structurally aligned with **GMe** (purple) co-crystallized DnaN (PDB: 8CIX).^12^ **(d – e)** Zoomed-in view of **GMe**-DnaN-α/ε interactions. Proteins are shown as surfaces; **GMe** is displayed as a space-filling model.

This renewed focus proved pivotal when researchers demonstrated that strategic structural modifications could dramatically enhance its therapeutic potential, transforming it into a promising anti-tubercular candidate. In 2015, *Kling et al.* reported new **GMe** derivatives by modifying Pro_8_ (the only non-methylated proline) with methyl and cyclohexyl moieties in the 4-position to create methyl griselimycin (**MeGMe**) and cyclohexyl griselimycin (**CyGMe**).^6^ **CyGMe**, possessing a larger and hydrophobic appendage from the Pro_8_ sidechain, in particular showed a marked increase in activity compared to **GMe**, improving the minimum inhibitory concentration (MIC) against *M. tuberculosis* from 0.9 µM to 0.05 µM. Interestingly, **CyGMe** displayed decreased binding affinity towards *M. tuberculosis* DnaN compared to **GMe** (K_D_ of 0.2 and 0.1 nM, respectively), suggesting that enhanced cellular accumulation of **CyGMe**, rather than improved target binding, could be driving its superior anti-tubercular activity. *In vitro* and *in vivo* degradation assays also showed that **CyGMe** was broadly more stable than **GMe**, which could contribute to its enhanced activity. Although the improved pharmacokinetics of **CyGMe** are promising, and it has been shown to be effective against both *M. tuberculosis* and *Mycobacterium abscessus* mouse infection models,^17^ no new derivatives have been reported since; **GMe** has also been explored as an adjuvant for anti-*M. tuberculosis* vaccines.^18^ A recent study of binding interactions between **GMe** and various DnaN homologues identified six positions with low binding pocket complementarity as potentially modifiable sites.^19^ While understanding clamp interactions and binding site architecture is useful for adapting **GMe** to target other organisms, the contribution of each residue towards activity in mycobacteria has yet to be explored.

To identify the residues necessary for activity and those amenable to modification, we performed an extensive structure-activity relationship (SAR) study of **GMe**. An alanine scan library of **GMe** analogs was synthesized and screened for antibiotic activity against *M. tuberculosis* and *Mycobacterium smegmatis*, a nonpathogenic model organism used to study *M. tuberculosis.*^20,21^ Other modifications previously suggested to impact peptide stability and permeability were assessed, including *N*-methylation of amino acids,^22,23^ replacement of the ester-based cyclization chemistry with an amide,^24,25^ and removal of the *N*-terminal acetyl group to expose a charged amine.^26^ Lastly, we made use of the SAR data to install an *N*-terminal fluorophore to enable visualization of griselimycin accumulation in *M. smegmatis* internalized within macrophage cells. From these results, we have identified positions Pro_2_ and Me-Val_7_ as promising sites of modification for designing future griselimycin derivatives.

## RESULTS AND DISCUSSION

### Assessment of Griselimycin Alanine Scan Library

To investigate the SAR of **GMe**, an alanine scan was performed (**Fig. 2a**). All peptides were synthesized using Fmoc-based solid-phase peptide synthesis (SPPS, **Scheme S1**). Derivatives were synthesized by sequentially replacing each amino acid with alanine, except for Thr_3_, which is necessary for cyclization (**Fig. 2a**). When replacing *N*-methylated positions (Val_1_, Val_7_, D-Leu_9_), variants were synthesized with both alanine and *N*-methyl-alanine while maintaining the stereochemistry; D-Leu_9_ was also replaced with L-alanine. In total, 13 alanine-scan derivatives were synthesized, accounting for all positions (except Thr_3_) and *N*-methylations.

**Figure 2.**
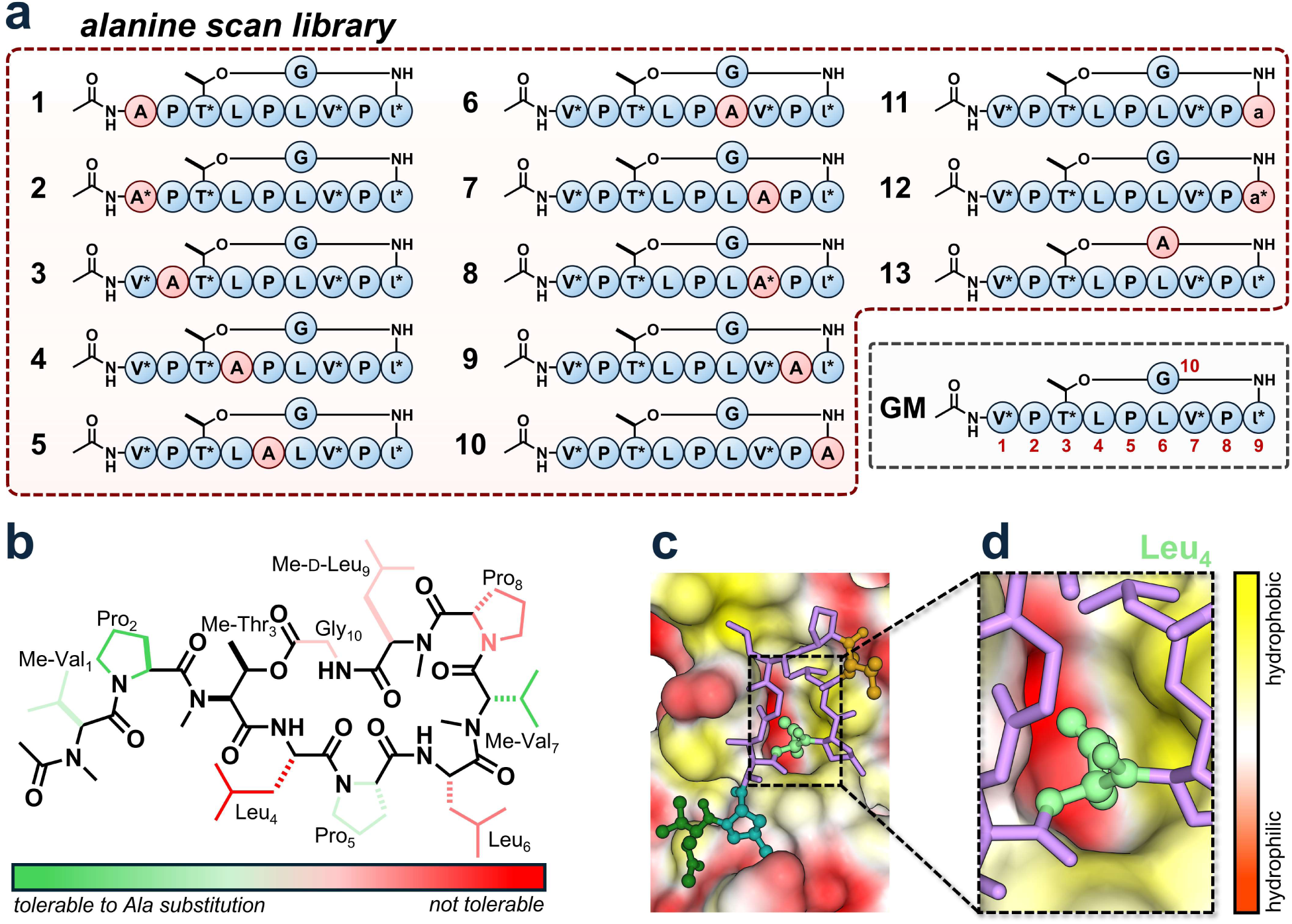
**(a)** Structures of **GM** and alanine derivatives. Asterisks (*) indicate N-methylated residues; residues in lowercase (only found in position 9) have D-stereochemistry. **(b)** Tolerability of **GM** residues to alanine substitution based on activity in *M. tuberculosis* MIC assays. Full data presented in **Table 1**. **(c)** Detail of **GM** binding pocket on DnaN (PDB: 5AGU)^6^ shown as a surface. Me-Val_1_, Pro_2_, Leu_4_, and Me-Val_7_ are highlighted in green, blue, purple, and gold respectively. **(d)** Zoomed-in view of the Leu4 binding pocket. Red-colored residues have hydrophilic side chains; yellow-colored residues have hydrophobic side chains.

**Table 1.**
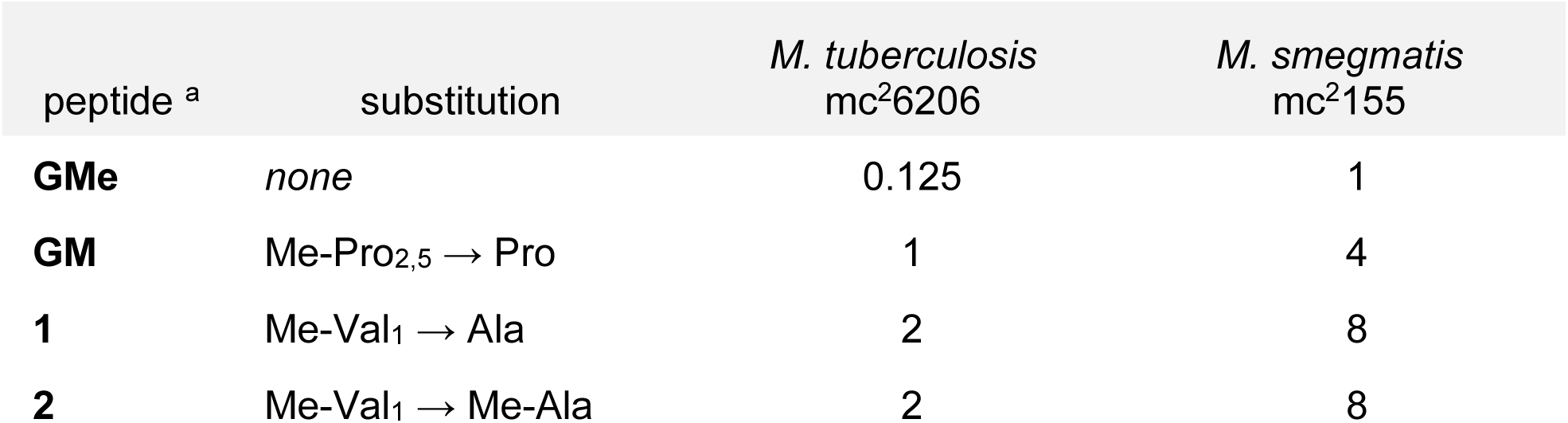
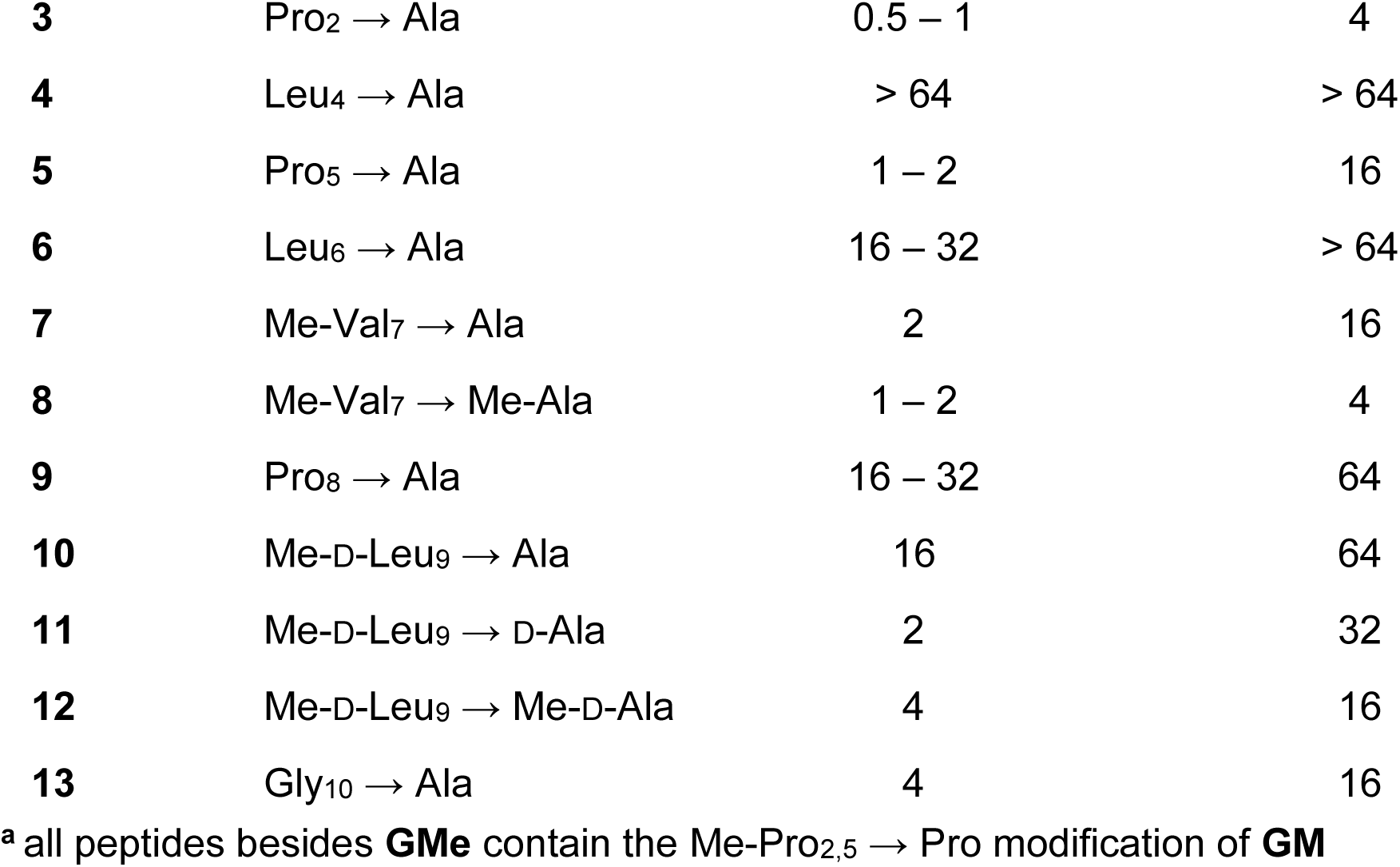
MICs (μM) of Alanine Scan Analogues of Griselimycin.

| peptide <sup>a</sup> | substitution | *M. tuberculosis*<br>mc<sup>2</sup>6206 | *M. smegmatis*<br>mc<sup>2</sup>155 |
| --- | --- | --- | --- |
| **GMe** | *none* | 0.125 | 1 |
| **GM** | Me-Pro<sub>2,5</sub> → Pro | 1 | 4 |
| **1** | Me-Val<sub>1</sub> → Ala | 2 | 8 |
| **2** | Me-Val<sub>1</sub> → Me-Ala | 2 | 8 |
| **3** | Pro<sub>2</sub> → Ala | 0.5 – 1 | 4 |
| **4** | Leu<sub>4</sub> → Ala | > 64 | > 64 |
| **5** | Pro<sub>5</sub> → Ala | 1 – 2 | 16 |
| **6** | Leu<sub>6</sub> → Ala | 16 – 32 | > 64 |
| **7** | Me-Val<sub>7</sub> → Ala | 2 | 16 |
| **8** | Me-Val<sub>7</sub> → Me-Ala | 1 – 2 | 4 |
| **9** | Pro<sub>8</sub> → Ala | 16 – 32 | 64 |
| **10** | Me-D-Leu<sub>9</sub> → Ala | 16 | 64 |
| **11** | Me-D-Leu<sub>9</sub> → D-Ala | 2 | 32 |
| **12** | Me-D-Leu<sub>9</sub> → Me-D-Ala | 4 | 16 |
| **13** | Gly<sub>10</sub> → Ala | 4 | 16 |

<sup>a</sup> all peptides besides **GMe** contain the Me-Pro<sub>2,5</sub> → Pro modification of **GM**

The natural product **GMe** contains two 4-methyl-prolines in positions 2 and 5 (**Fig. 1a**). Due to the difficulty in synthetic accessibility to the building block Fmoc-4-methyl-proline and our large library size, all derivatives in this study were synthesized using standard unmethylated proline (**Fig. 2a**). To ensure the 4-methyl-prolines were not essential for activity, **GMe** was purchased and tested against a nonmethylated version, **GM**, (**Fig. 1a**) *via* minimum inhibitory concentration (MIC) assay (**Fig. 2b**; **Table 1**). **GM** had a four to eightfold higher MIC than **GMe** against both *M. tuberculosis* and *M. smegmatis* (1 and 4 µM, respectively), suggesting that methylation of Pro_2_ and Pro_5_ plays an important but nonessential role in the antibiotic activity of the peptide.

Alanine-substituted griselimycins displayed a range of activities against mycobacteria (**Fig. 2b**; **Table 1**). Interestingly, both exocyclic residues (Me-Val_1_ and Pro_2_) had MICs equivalent or close to **GM** upon alanine substitution (compounds **2** and **3**, respectively). These sites are expected to exhibit higher inherent flexibility and thus contribute less to the binding interaction with DnaN, consistent with observations from the crystal structure. The high tolerability of Me-Val_1_, Pro_2_, and Me-Val_7_ to alanine substitution may be due to the solvent-facing position of their side chains as observed in **GMe**-DnaN crystal structures (**Fig. 2c**). Of the macrocyclic residues, Me-Val_7_ was the least affected by alanine substitution (**8**), displaying the same activity as **GM** against both *M. tuberculosis* and *M. smegmatis*. Pro_5_ substitution was well tolerated in *M. tuberculosis* but showed poorer activity in *M. smegmatis* (**5**), while Me-D-Leu_9_ substitution resulted in a fourfold increase in MIC against both organisms (**12**). Leu_6_ and Pro_8_ substitution sharply raised the MIC to 16 – 32 µM in *M. tuberculosis* (**6** and **9**), whereas Leu_4_ was the only position for which no growth inhibition was observed against *M. tuberculosis* or *M. smegmatis* at concentrations ≤ 64 µM (**4**). The side chain of Leu_4_ is buried deep within an amphipathic pocket on DnaN and appears to make the most contact with the protein of all **GMe** residues, which may explain the complete lack of activity upon alanine substitution (**Fig. 2c - d**). This is corroborated by a recent study demonstrating that alanine substitution at Leu_4_ reduced interaction with *M. smegmatis* DnaN by approximately 100-fold relative to **GM**, as measured by competitive fluorescence polarization assays.^27^ Overall, changes in MIC upon alanine substitution were mostly consistent between *M. tuberculosis* and *M. smegmatis*, with Pro_2_ and Leu_4_ being the positions most and least tolerable to alanine substitution, respectively. These findings suggest that the nonpathogenic model organism *M. smegmatis* could serve as a viable surrogate for evaluating advanced griselimycin analogs in future studies.

Maintaining the *N*-methylation of Me-Val_7_ in the alanine substitution analogue **8** afforded a slight increase in activity compared to an unmethylated version (**7**), whereas demethylating Me-D-Leu_9_ (**11**) slightly improved activity against *M. tuberculosis*. (**Table 1**). The configuration of Leu_9_ had a strong effect on activity towards *M. tuberculosis*, with an eightfold reduction in MIC observed upon exchanging D-alanine (**11**) for L-alanine (**10**, **Table 1**). Me-Val_1_ had identical activity when converted to alanine (**1**) or *N-*methyl-alanine (**2**). While an alanine-substitution at Me-Thr_3_ could not be assessed, a non-*N*-methylated threonine analogue, **14**, was tested (**Fig. 3a**); absence of *N*-methylation at this position did not affect the activity against *M. smegmatis* and only slightly increased the MIC against *M. tuberculosis* (**Table 2**). Although not tested here, previous work by our group has shown that removing all four *N*-methylations in **GM** eliminates antibiotic activity^28^; thus the contributions of each *N*-methylation may be additive.

**Figure 3.**
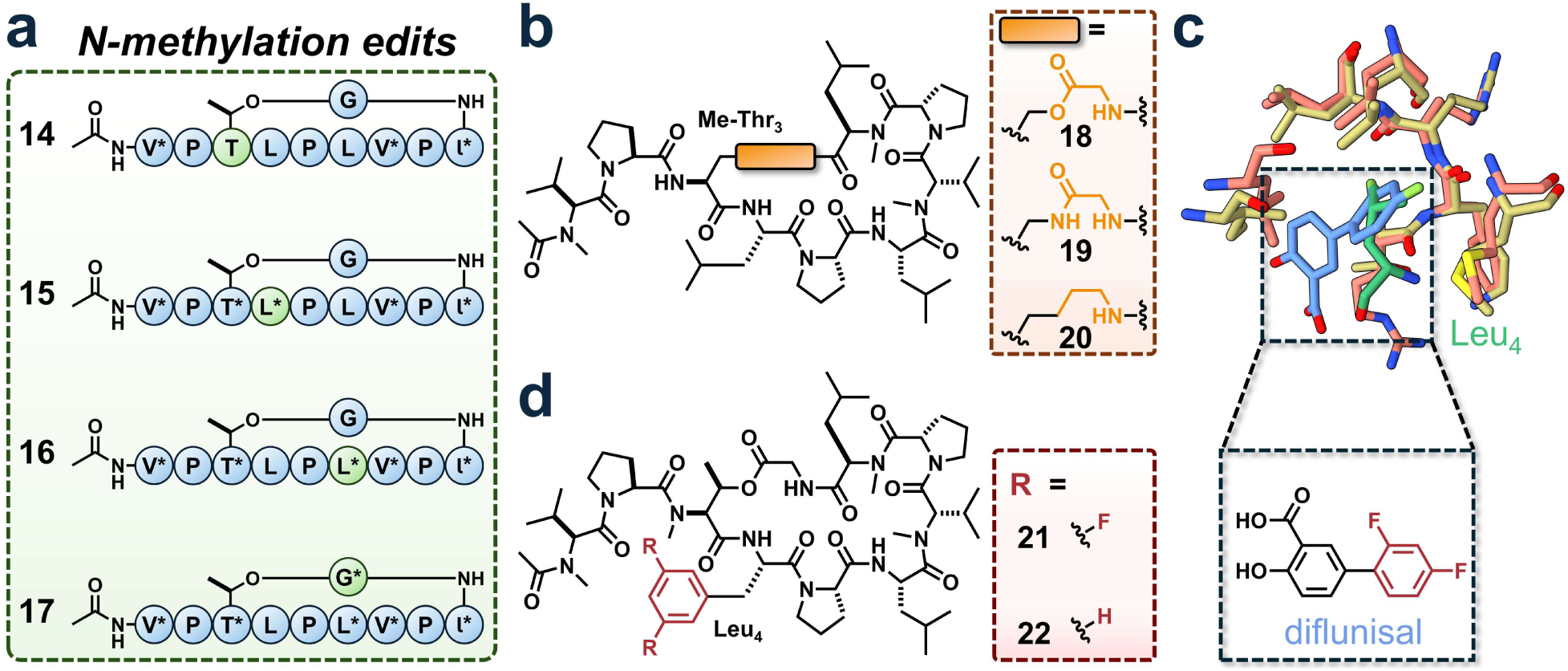
**(a)** Structures of GM derivatives with removed (**14**) or added (**15**, **16**, **17**) *N*-methylation. **(b)** Structures of GM derivatives with edits to Thr_3_ (**18**, **19**, **20**). **(c)** Detail of Leu_4_ from GM (green) and diflunisal (blue) bound to *M. tuberculosis* (red) and *H. pylori* (tan) DnaN respectively (PDBs: 5AHU and 5G48).^6,29^ The difluorophenyl ring of diflunisal is highlighted in red. **(d)** Structures of GM derivatives with edits to Leu_4_ (**21**, **22**). MIC values are shown in **Table 2**.

**Table 2.** MICs (μM) of *N*-methyl, Side Chain, and Cyclization-Modified Derivatives of Griselimycin.

| peptide <sup>a</sup> | substitution | *M. tuberculosis*<br>mc<sup>2</sup>6206 | *M. smegmatis*<br>mc<sup>2</sup>155 |
| --- | --- | --- | --- |
| **14** | Me-Thr<sub>3</sub> → Thr | 2 | 4 |
| **15** | Leu<sub>4</sub> → Me-Leu | > 64 | > 64 |
| **16** | Leu<sub>6</sub> → Me-Leu | > 64 | > 64 |
| **17** | Gly<sub>10</sub> → Me-Gly | > 64 | > 64 |
| **18** | Me-Thr<sub>3</sub> → Ser | 4 – 8 | 16 |
| **19** | Me-Thr<sub>3</sub> → Dap <sup>b</sup> | 64 | > 64 |
| **20** | Me-Thr<sub>3</sub> and Gly<sub>10</sub> → Lys | > 32 | > 32 |
| **21** | Leu<sub>4</sub> → DiF-Phe <sup>c</sup> | 32 | 64 |
| **22** | Leu<sub>4</sub> → Phe | 8 | 16 |

<sup>a</sup> all peptides contain the Me-Pro<sub>2,5</sub> → Pro modification of **GM**
<sup>b</sup> Dap = 2,3-diaminopropionic acid
<sup>c</sup> DiF-Phe = 3,5-difluorophenylalanine

As **GMe** contains four *N*-methylated residues and three prolines, only three positions (Leu_4_, Leu_6_, and Gly_10_) have backbone secondary amides. The prevalence of alkylated backbone positions, whether through *N*-methylation or proline incorporation, may enhance passive membrane permeability by reducing hydrogen bonding potential and increasing lipophilicity of the peptide. To examine how adding *N*-methylation at these positions affects activity, three additional derivatives of **GM** were synthesized (**15**, **16**, and **17**; **Fig. 3a**). All three compounds displayed a MIC > 64 µM against both *M. tuberculosis* and *M. smegmatis* (**Table 2**). Interestingly, nearly all residues with a backbone nitrogen within the cyclic portion of the peptide showed a decrease in activity when changing the native *N*-methylation state, while the exocyclic Me-Val_1_ was unperturbed. As addition or removal of an *N*-methyl group changes the number of hydrogen bond donors, this observation may be a consequence of disrupting an optimal intramolecular hydrogen bonding network within the macrocycle of **GM** which is necessary for activity.

### Substitution of the Thr_3_-Gly_10_ Ester for an Amide

Mycobacteria are known to express esterases,^30,31^ which could hydrolyze the ester bond between the side chain of Me-Thr_3_ and Gly_10_ to linearize griselimycin and reduce DnaN affinity. This ester linkage could also represent a liability in human serum or within host macrophages, where esterase activity may prematurely hydrolyze the cyclic peptide and generate an inactive linear metabolite before the compound reaches the mycobacterial target. Converting this ester to a more metabolically stable functional group could maintain the cyclic form of the peptide and potentially increase activity. To test this, a variant of **GM** with an amide instead of an ester (**19**) was synthesized by replacing Me-Thr_3_ with an amine-containing mimic, 2,3-diaminopropionic acid (Dap; **Fig. 3b**). This substitution was expected to retain the cyclic structure and binding conformation while eliminating susceptibility to esterase-mediated hydrolysis. The lack of synthetic accessibility of the corresponding Dap building block, containing both backbone *N*-methylation and a β-methyl substituent, led to the evaluation of another **GM** derivative substituting serine for Me-Thr_3_ (**18**; **Fig. 3b**).

Interestingly, our data showed that **19** exhibited no antibiotic activity against *M. tuberculosis* or *M. smegmatis* at concentrations ≤ 64 µM (**Table 2**). Conversely, **18** showed four to eightfold reduced activity towards *M. tuberculosis* compared to **GM**. As **18** and **19** differ only in the substitution of the ester functional group for an amide, presence of an ester at this position appears necessary for proper activity. Removal of the ester functional group was further assessed by substituting both Me-Thr_3_ and Gly_10_ with lysine and cyclizing via amide bond formation between the amino side chain and the C-terminal Me-D-Leu_9_ (**20**; **Fig. 3b**). This modification maintains the same number of atoms in the cyclic backbone as **GM** while replacing the ester with two methylene groups. As with **19**, **20** displayed no antibiotic activity against *M. smegmatis* or *M. tuberculosis*, underscoring the necessity of the Thr_3_-Gly_10_ ester (**Table 2**). Similar phenomena have been observed when replacing esters with amides in natural compounds including small molecules and other threonine-cyclized peptides.^32,33^

It is possible that the double bond character of the amide N–C bond disrupts **19** from achieving ideal geometry for DnaN engagement, although crystallography and/or binding affinity assays would be necessary to confirm this. Alternatively, in a tightly constrained cyclic peptide, introducing a new N–H donor into the backbone alters the global energy landscape. The macrocycle will typically contort to satisfy this new hydrogen bond donor, forging a novel intramolecular hydrogen bond (IMHB) network. This thermodynamic reorganization often collapses the peptide into an entirely different, tightly folded ground-state conformation that misaligns the critical side chains required for engaging the target (like DnaN).^34^ Finally, the energetic penalty for stripping water away from an exposed, polar amide N–H is significantly higher than the penalty for desolvating an ester oxygen.^24^ As the non-*N*-methylated **14** displayed only slightly reduced MIC compared to **GM**, the β-methyl group absent from **19** appears to be important but nonessential for activity. **GMe**-bound crystal structures of DnaN have this methyl group poised in an amphipathic pocket on the surface of the protein, where it may influence binding. However, as with all other changes to **GM**, numerous facets such as membrane permeability, metabolic stability, or DnaN binding may be affected in complementary or opposite ways.

The effects of cyclization chemistry on the stability of **GM** and **19** were assessed in both human serum and mycobacterial cell lysate via time course degradation assays. We observed a slightly higher rate of degradation for **GM** compared to **19** in human serum (**Fig. S1**), which may be due to the presence of hydrolytic esterase enzymes.^35^ Interestingly, this trend was reversed in *M. smegmatis* cell lysate, where the concentration of **19** nearly halved by 30 minutes while **GM** appeared unaffected (**Fig. S1**). Although mycobacteria are known to express numerous esterases,^30,36^ it is possible that a privileged hydrogen bonding state or higher DnaN occupancy affords **GM** greater resistance to degradation by mycobacterial lysate, which may explain the lack of any notable growth inhibition by **19** in MIC assays (**Table 2**).

### Incorporation of an Unnatural Amino Acid at Leu_4_

Unnatural amino acids are a useful source of building blocks for derivatizing peptide therapeutics as they enable natural products to sample chemical spaces inaccessible to living organisms.^37^ Numerous small molecule inhibitors of DnaN have been described,^38^ providing a variety of chemical motifs to mimic on griselimycin. Of note is the salicylic acid derivative diflunisal, which has been shown to inhibit DnaN function in *Helicobacter pylori*.^29^ Structure alignments of deposited *H. pylori* and *M. tuberculosis* DnaN crystal structures suggest that diflunisal bind by burying its difluorophenyl ring into the same conserved pocket as Leu_4_ (**Fig. 3c**).^6,29^ As Leu_4_ was shown to be the position least tolerable to alanine substitution and is likely a key DnaN binding residue, a derivative of **GM** was synthesized replacing Leu_4_ with 3,5-difluorophenylalanine (**21**) to mimic diflunisal and potentially increase clamp affinity (**Fig. 3d**). An equivalent phenylalanine derivative (**22**) was also synthesized to assess the contributions of ring size and fluorination (**Fig. 3d**).

**22** was found to have poor activity in both *M. tuberculosis* and *M. smegmatis*, with eight- and fourfold-increased MIC relative to **GM** (**Table 2**). Activity of **21** was further compromised with MICs of 32 and 64 µM, analogous to the activity of diflunisal against *H. pylori* (MIC of 84 µM).^29^ As *H. pylori* and *M. smegmatis* DnaN only show moderate sequence similarity at the **GM** binding site (strict conservation at 11 of 21 residues), differences in protein architecture may explain the poor performance of these derivatives (**Fig. S2**); the Leu_4_/diflunisal binding pocket, however, appears to be conserved between *H. pylori* and *M. smegmatis* DnaN (**Fig. 3b**). It is possible that the difluorophenyl-DnaN interaction in diflunisal is weaker than the analogous Leu_4_-DnaN interaction in **GM**. The additional size and planarity of the phenylalanine ring relative to leucine may also disrupt binding, as evidenced by the poor activity of **22**. The sharp reduction of activity observed in **21** and **22** suggests the size and conformational flexibility afforded by Leu_4_ is critical for proper engagement of DnaN by **GM**, although additional binding affinity experiments may be required to confirm this.

### Proline Ring and Side Chain Modifications

Noting the tolerability of Pro_2_ and Pro_5_ to alanine substitution and the increased antitubercular activity of Pro_8_-modified **MeGMe** and **CyGMe**,^6^ we assessed the effects of various proline modifications and substitutions. We synthesized three derivatives of **GM** by substituting L-homoproline for Pro_2_, Pro_5_, and Pro_8_ (**23**, **24**, and **25**, respectively), surmising that a nonproteinogenic amino acid may reduce the susceptibility of **GM** to mycobacterial proteases (**Fig. 4a**). Compared to **GM**, **24** displayed twofold improvements in MIC against both *M. smegmatis* and *M. tuberculosis*, while the activity of **25** was unchanged (**Table 3**). **23** was not tested for antibacterial activity as the ester appeared to be hydrolyzed under HPLC conditions, which may indicate an unfavorable change in the secondary structure. Both *M. smegmatis* lysate and human serum time course degradation assays indicated that the stability of **24** and **25** was relatively unchanged compared to **GM**, with all three peptides displaying ≤ 15% degradation after 1-hour incubations (**Fig. S3**). Therefore, the additional ring size afforded by homoproline substitution at Pro_5_ may alter the backbone conformation of **24** to favor increased permeability or target affinity.

**Figure 4.**
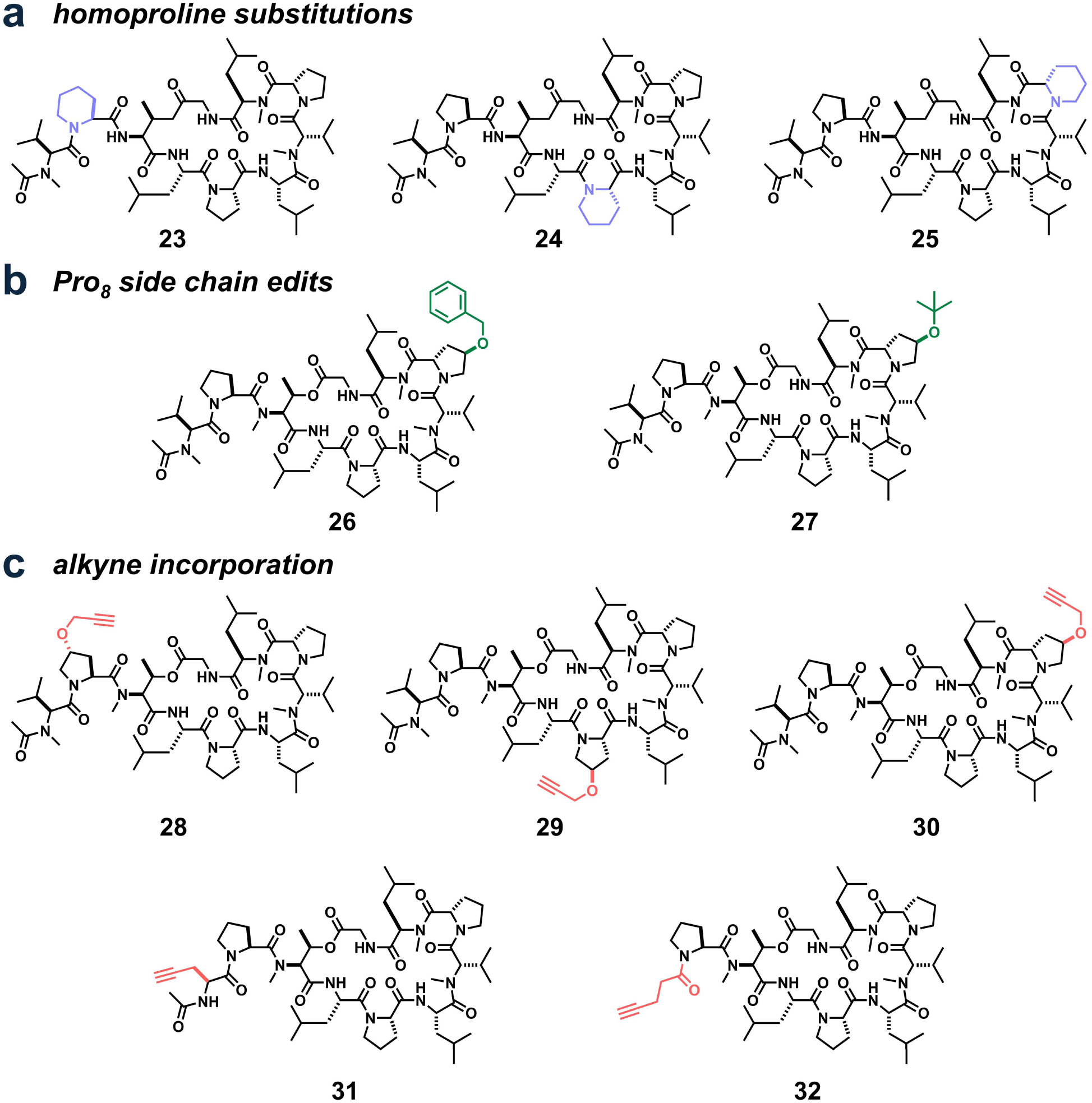
**(a)** Structures of homoproline-substituted griselimycins. **(b)** Structures of Pro_8_ side chain modified griselimycins. **(c)** Structures of alkynylated griselimycins. MIC values are shown in **Table 3**.

**Table 3.** MICs (μM) of Modified Proline and Alkyne Derivatives of Griselimycin.

| peptide <sup>a</sup> | substitution | *M. tuberculosis*<br>mc<sup>2</sup>6206 | *M. smegmatis*<br>mc<sup>2</sup>155 |
| --- | --- | --- | --- |
| **23** | Pro<sub>2</sub> → HomoPro | N/A | N/A |
| **24** | Pro<sub>5</sub> → HomoPro | 0.5 | 2 |
| **25** | Pro<sub>8</sub> → HomoPro | 1 | 4 |
| **26** | Pro<sub>8</sub> → 4-benzyloxy-Pro | 0.25 | 1 |
| **27** | Pro<sub>8</sub> → 4-*tert*-butoxy-Pro | 0.25 | 1 |
| **28** | Pro<sub>2</sub> → 4-propynyloxy-Pro | 0.25 | 1 |
| **29** | Pro<sub>5</sub> → 4-propynyloxy-Pro | 0.5 | 4 |
| **30** | Pro<sub>8</sub> → 4-propynyloxy-Pro | 0.25 – 0.5 | 1 |
| **31** | Me-Val<sub>1</sub> → Pra | 1 | 16 |
| **32** | Me-Val<sub>1</sub> → 4-pentynone | 1 | 8 |

<sup>a</sup> unless modified otherwise, all peptides contain the Me-Pro<sub>2,5</sub> → Pro modification of **GM**

Although Pro_8_ did not tolerate alanine substitution, the improved activities of **MeGMe** and **CyGMe** suggest that derivatization off the 4-position of the ring is a viable route to improve griselimycin activity. To further assess this, we synthesized two variants of **GM** incorporating either benzyl (**26**) or *tert*-butyl ether (**27**) moieties on the 4-position of Pro_8_ (**Fig. 4b**). Both derivatives displayed fourfold improvement in MIC over **GM** in both *M. smegmatis* and *M. tuberculosis*. As **MeGMe** and **CyGMe** were not shown to increase DnaN binding affinity,^6^ it is plausible that the increased activity of **26** and **27** is a consequence of other mechanisms.

### Alkyne Griselimycin Scaffolds

The copper-catalyzed azide-alkyne cycloaddition (CuAAC) is widely used in drug discovery for rapid and facile diversification of small molecules.^39,40^ The location of a click chemistry handle on a molecule can affect its physiochemical properties and is therefore of careful consideration during drug development. To assess the viability of griselimycin as a CuAAC scaffold, we synthesized five alkyne-containing derivatives of **GM** (**Fig. 4c**). As we have shown the 4-position of the Pro_8_ ring as a viable site of functionalization, we substituted an ether-linked alkyne at this position for Pro_2_ (**28**), Pro_5_ (**29**), and Pro_8_ (**30**). Due to the exocyclic position and strong performance of Me-Val_1_ in the alanine scan, we also synthesized two alkyne substitutions at this position (**31** and **32**).

MIC assays showed that alkyne incorporation on **GM** gives location-dependent changes in activity (**Table 3**). Addition of an alkyne on side chains of Pro_2_ (**28**) and Pro_8_ (**30**) afforded a fourfold decrease in MIC compared to **GM**, whereas alkynylation at Pro_5_ gave only a twofold increase in activity against *M. tuberculosis* and no increase in activity against *M. smegmatis* (**29**). *N*-terminal alkynes gave no change in activity against *M. tuberculosis* compared to **GM**, but faired worse in *M. smegmatis* with four- and twofold increases in MIC (**31** and **32**). In conjunction with other Pro_8_ edits (**25** and **26**), the strong performance of **30** underscores the potential for side chain expansion at Pro_8_ to improve activity.

### N-terminal Modifications to Griselimycin

Given the strong tolerance of exocyclic residues Me-Val_1_ and Pro_2_ to alanine substitution and its potential for late-stage diversification after cyclization, the *N*-terminus of **GMe** was examined as a site of modification. While **GMe** and all derivatives in this study contain an acetylated *N*-terminus, the presence of a primary amine on small molecules has been associated with enhanced membrane permeability in Gram-negative bacteria.^26,41^ To examine whether this modification could lead to improve activity in mycobacteria, derivatives of **GM** were synthesized with a free *N*-terminal primary amine (**33**) and an appended ornithine residue (**34**; **Fig. 5a**). Noting the identical activity of **1** and **2**, the *N*-methylation normally present at Val_1_ was omitted to generate a primary amine as opposed to a secondary amine for **33**. **33** displayed the same activity as **GM** in *M. smegmatis*, while MIC was reduced to 4 µM against *M. tuberculosis* (**Table 4**). Like other derivatives, it is possible that free amine improved activity in one aspect, such as increased accumulation, while reducing it in another. While crystal structures do not reveal any obvious interactions of the acetyl group with DnaN, the free *N*-terminus could experience repulsion with neighboring cationic residues in the binding pocket and reduce affinity (**Fig. 2c**). As the *C*-terminus of **GM** is masked by cyclization, acetylation of the *N*-terminus may help protect the exposed end of the peptide from exopeptidase activity.^42^ Consequentially, removing the acetyl group may make **33** more susceptible to degradation. By contrast, addition of an *N*-terminal ornithine in compound **34** (thereby also adding a free primary amine) resulted in a twofold improvement in MIC against both *M. smegmatis* and *M. tuberculosis* compared to **GM** (**Table 4**).

**Figure 5.**
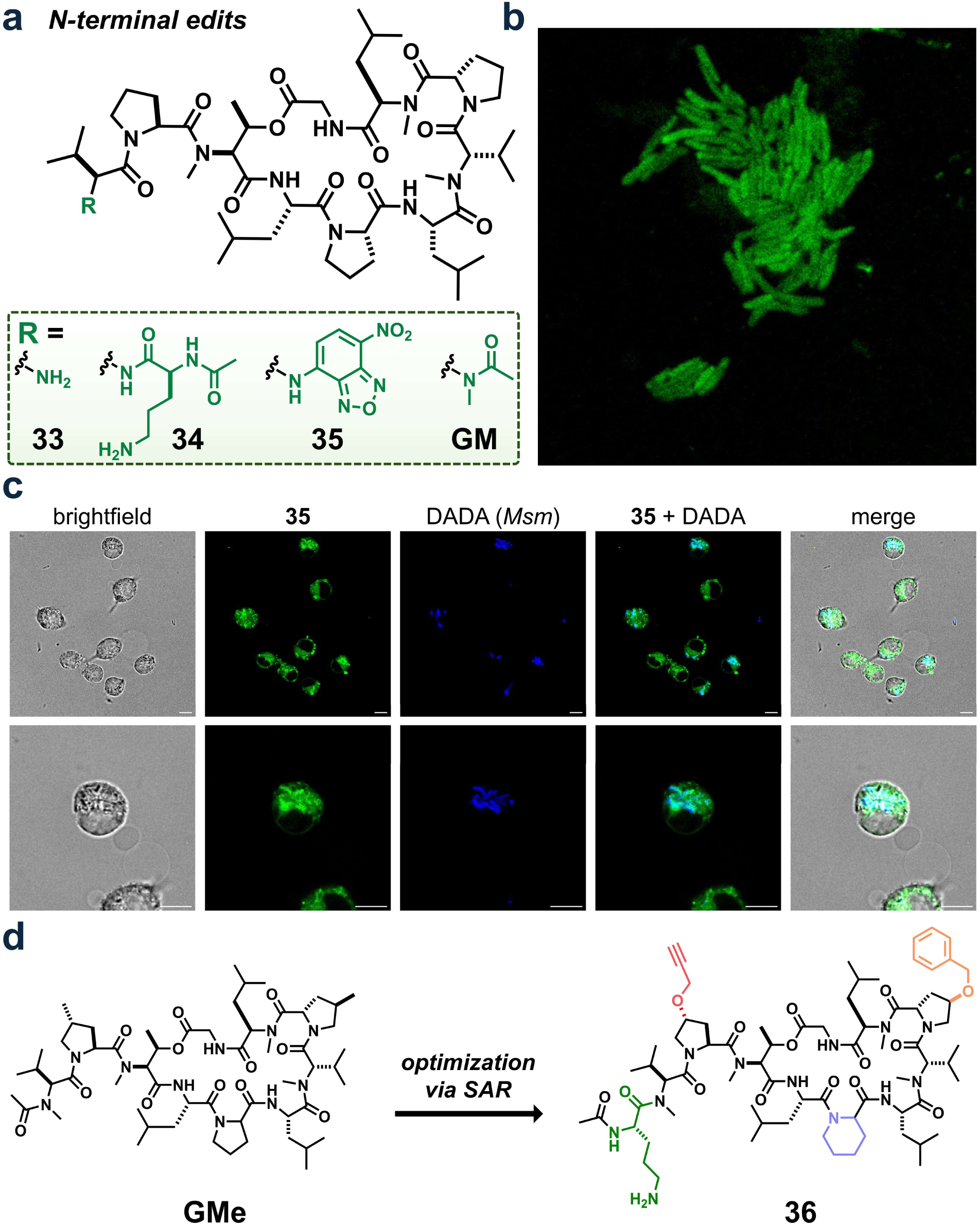
**(a)** Structures of *N*-terminal derivatives of **GM**. MIC values are shown in **Table 4**. **(b)** Confocal analysis of *M. smegmatis* incubated with 10 µM of **35**. **(c)** Confocal analysis of PG-labeled *M. smegmatis* (DADA, 100 µM) in J774A.1 macrophages treated with **35** (10 µM). Scale bars represent 10 µm. **(d)** Structure of **GMe** derivative **36**, incorporating multiple edits individually shown to increase activity.

**Table 4.** MICs (μM) of *N*-terminal derivatives of Griselimycin.

| peptide <sup>a</sup> | substitution(s) | *M. tuberculosis*<br>mc<sup>2</sup>6206 | *M. smegmatis*<br>mc<sup>2</sup>155 |
| --- | --- | --- | --- |
| **33** | *N*-terminal NH<sub>2</sub> | 4 | 4 |
| **34** | *N*-terminal Orn | 0.5 | 2 |
| **35** | *N*-terminal NBD <sup>b</sup> | 0.5 – 1 | 2 |
| **36** | *N*-terminal Orn<br>Pro<sub>2</sub> → 4-propargyloxy-Pro<br>Pro<sub>5</sub> → HomoPro<br>Pro<sub>8</sub> → 4-benzyloxy-Pro | 0.25 | 0.5 – 0.25 |

<sup>a</sup> peptides **33**, **34**, and **35** contain the Me-Pro<sub>2,5</sub> → Pro modification of **GM**
<sup>b</sup> NBD = nitrobenzofurazan

To visualize the accumulation of **GM** in *M. smegmatis*, a variant was synthesized with the small molecule fluorophore nitrobenzofurazan (NBD) appended to the *N*-terminus (**35**, **Fig. 5a**). NBD was selected due to its small size, which minimizes perturbation of the peptide’s physicochemical properties, and its environment-sensitive fluorescence that allows discrimination between intracellular and extracellular localization.^43^ Confocal imaging revealed that **35** produced a uniform fluorescent signal throughout *M. smegmatis* when incubated at 10 µM, suggesting effective uptake and distribution (**Fig. 5b**). Interestingly, the MIC of **35** was slightly improved compared to **GM** in both *M. tuberculosis* and *M. smegmatis*. Along with **33** and **34**, this suggests that the *N*-terminus of **GM** is flexible to modification and may represent a promising site for late-stage derivatization.

### Activity of Griselimycins Against Escherichia coli

As the mycobacterial membrane can present a significant accumulation barrier for cytosolically active antibiotics,^44^ we sought to assess the activity of griselimycins in Gram-negative bacteria. The cellular envelope of Gram-negative bacteria features a thin PG layer sandwiched between compositionally distinct inner and outer membranes, whereas mycobacteria have a single inner membrane surrounded by a larger PG and a thick layer of mycolic acids.^45^ Interestingly, we found that **GM** had no measurable effect on the growth of *Escherichia coli* at concentrations ≤ 32 μM (**Fig. S4**). Noting that the lack of **GM** activity in *E. coli* may be a consequence of poor accumulation due to differences in cell envelope architecture, we also assessed the effect of polymyxin B nonapeptide (PMBN) coincubation. PMBN is known to permeabilize the outer membrane of Gram-negative bacteria without significant bactericidal activity.^46^ Accordingly, if accumulation is the bottleneck for **GM** activity in Gram-negative bacteria, we may expect an improved MIC in *E. coli* upon PMBN coincubation.

We found that *E. coli* growth was unperturbed in the presence of **GM** and was unaffected by PMBN coincubation (**Fig. S4**). Interestingly, the primary amine-containing derivatives **33** and **34** also displayed no activity against *E. coli* with or without PMBN coincubation, despite the reported enhanced permeability of such molecules in Gram-negative bacteria (**Fig. S4**).^26^ This may indicate that membrane permeability is not the limiting factor for **GM** activity in Gram-negative bacteria. Indeed, **GM** is reported to bind *E. coli* DnaN with 8000-fold poorer affinity than *M. smegmatis* DnaN (K_D_ of 650 and 0.08 nM, respectively), and in general shows poor activity against Gram-negative organisms.^6^ These results corroborate with other published literature, which showed that **GMe** has significantly lower affinity for non-mycobacterial DnaN.^19^ Notably, *M. smegmatis* DnaN has poorer sequence similarity with *E. coli* (71% similarity; strict conservation at 14 of 21 binding pocket residues) compared to *M. tuberculosis* (92% similarity; strict conservation at 20 of 21 binding pocket residues; **Fig. S2**). This suggests that strong homology with the *M. tuberculosis* and *M. smegmatis* DnaN binding pockets is necessary for sufficient **GM** activity against a given bacterium.

### Assessment of Griselimycin Accumulation in Infected Macrophages

*M. tuberculosis* replicates and resides within phagosomes of host macrophages.^47,48^ This presents two additional permeability barriers, the macrophage and phagosomal phospholipid bilayers, which *M. tuberculosis*-targeting drugs must cross before reaching the cytosol. To determine if griselimycin derivatives can access mycobacterial cells within macrophages, we utilized a fluorescent D-alanine analogue (DADA) which is incorporated into the peptidoglycan (PG) of mycobacteria (**Fig. S5**).^49^ The peptidoglycan layer of *M. smegmatis* was metabolically labeled with DADA prior to the infection of J774A.1 macrophages, which were subsequently incubated with **35**. Confocal microscopy revealed the DADA fluorescent signal to be localized within the phagosome and visually corresponding with intact bacteria (**Fig. 5c**). **35** fluorescence strongly colocalized with the phagocytosed DADA-labeled bacteria, indicating that griselimycin can access mycobacteria within macrophages (**Fig. 5c**). This finding suggests that griselimycin possesses the necessary physicochemical properties to traverse multiple biological barriers and reach its intracellular target, and corroborates previously published intramacrophage activity studies.^6^ The successful accumulation within the phagosomal environment represents a critical pharmacological advantage for treating TB infections in their physiologically relevant host cell context.

### Combined Modifications to Griselimycin Increase Potency

To assess if the effects of various modifications were additive, we synthesized a variant of **GMe** containing four edits shown to individually increase antibiotic activity (**36**). Compared to the natural product **GMe**, this peptide contains: (i) an *N*-terminal ornithine residue; (ii) a 4-propynyloxy-proline substitution at Pro_2_; (iii) a homoproline substitution at Pro_5_; and a 4-benzyloxy-proline substitution at Pro_8_ (**Fig. 5d**). Individually, these modifications afforded two-to fourfold decreases in MIC against *M. smegmatis* and *M. tuberculosis*. Remarkably, their effects appear to be additive, with a MIC of 0.5 – 0.25 μM against *M. smegmatis* representing an eight-to sixteenfold improvement in activity over **GM** and surpassing that of the natural product **GMe** (**Table 4**). A strong MIC of 0.25 μM was observed against *M. tuberculosis* for **36**, although this was twofold higher than **GMe**. The potential for increased activity by diversifying via the alkyne handle installed at Pro_2_ makes **36** a promising design for further antimycobacterial development.

## CONCLUSION

In this work, we present a comprehensive SAR study of the antitubercular cyclic peptide griselimycin. The unnatural side chain methylation of Pro_2_ and Pro_5_ is important but nonessential for activity. An alanine scan revealed that the exocyclic residue Pro_2_ is the most tolerable site to alanine substitution, whereas most macrocyclic residues gave strong decreases in activity upon change. Among these, Leu_4_ was the least tolerable to modification, consistent with its side chain residing deep within a hydrophobic pocket of DnaN.

Backbone *N*-methylation exhibited position-dependent effects. Macrocyclic residues generally showed a decrease in activity upon removal of native methylation, whereas demethylation of exocyclic residues had minimal effect. In contrast, additional *N*-methylation at the three native secondary amides abolished activity, suggesting that the *N*-methylation state of the macrocycle is finely tuned. This arrangement likely optimizes intramolecular hydrogen bonding and/or interactions within the DnaN binding pocket, a conclusion further supported by the loss of activity upon replacing the cyclizing ester bond for an amide, which may disrupt the conformational flexibility.

Several proline modifications with improved MIC were identified, including ring expansion and Pro_8_ side chain functionalization. Alkynylation gave location-dependent changes in activity, with Pro_2_ and Pro_8_ alkynes being potential scaffolds for CuAAC derivatization. We showed that a fluorophore-labeled griselimycin analogue was spatially coincident with mycobacteria located within macrophage phagosomes, highlighting its potential as a therapeutic agent. A final design combining the most productive edits, compound **36**, showed improved activity compared to our parent structure and can be further diversified using click chemistry. Overall, this SAR provides mechanistic insights into the function of griselimycin and establishes a foundation for the rational design of future analogs optimized for antimycobacterial activity.

## EXPERIMENTAL

### Synthesis of alanine scan griselimycin library

All peptides were synthesized via Fmoc-based solid phase peptide synthesis using a previously reported route (**Scheme S1**).^50^ Briefly, 2-chlorotritylchloride resin (1 eq., 0.25 mmol, 1.55 mmol/g loading capacity) was added to an oven-dried peptide synthesis vessel. Fmoc-*N*-methyl-D-leucine (1.1 eq., 0.275 mmol) was dissolved in anhydrous DCM (5 mL), to which anhydrous DIEA (4 eq, 1 mmol) was added. This solution was added to the resin and shaken at room temperature for 1 h. The resin was washed (∼5 mL each of DCM, MeOH, DCM, MeOH, DCM, DCM) and the Fmoc protecting group was removed by shaking with 10 mL of piperidine in DMF (20% v/v) for 30 minutes then washing as described. Each sequential Fmoc-protected amino acid (5 eq., 1.25 mmol) was coupled by adding a solution of Oxyma (5 eq., 1.25 mmol) and DIC (5 eq., 1.25 mmol) in DMF (7 mL) to the resin, shaking for 2 h at room temperature, then washing and deprotecting as described. The sequence of **GM** was followed, replacing the appropriate amino acid with Fmoc-*N*-methyl-alanine-OH (**2**, **8**), Fmoc-D-*N*-methyl-alanine-OH (**11**), or Fmoc-alanine-OH (all others). To acetylate the *N*-terminus, acetic acid (5 eq., 1.25 mmol) was coupled in the same manner as above. The peptide was esterified via Steglich esterification by adding Fmoc-glycine (10 eq., 2.5 mmol), DIC (10 eq., 2.5 mmol), and DMAP (0.25 eq., 0.063 mmol) dissolved in anhydrous THF (10 mL) to the resin and shaking at room temperature overnight. In the case of **13**, Fmoc-alanine-OH was used instead. The Fmoc protecting group was then removed as described. Peptides were cleaved from the resin by adding a solution of HFIP in DCM (30% v/v containing 2.5% v/v triisopropylsilane; 10 mL total) and shaking at room temperature for 1 h. This solution was filtered, concentrated, and washed with hexanes (3 x 5 mL) followed by a 1:1 solution of hexanes in cold diethyl ether (10 mL). The peptide was dissolved in ACN (15 mL) and cyclized by adding this solution dropwise to a solution of COMU (3 eq., 0.75 mmol) and DIEA (6 eq., 1.5 mmol) dissolved in ACN (150 mL) and stirring at room temperature overnight. This solution was concentrated and washed with hexanes (3 x 5 mL) followed by a 1:1 solution of hexanes in cold diethyl ether (10 mL) to afford crude peptide.

#### Synthesis of 14, 15, 16, 17, 18, 21, and 22

Peptides were synthesized as described above for **GM**, replacing the appropriate amino acid with either Fmoc-threonine-OH (**14**), Fmoc-*N*-methyl-leucine-OH (**15** and **16**), Fmoc-sarcosine-OH (**17**), Fmoc-serine-OH (**18**), Fmoc-phenylanaine(3,5-difluoro)-OH (**21**), or Fmoc-phenylalanine-OH (**22**).

#### Synthesis of 19

Fmoc-Gly (1.1 eq., 0.275 mmol) was coupled to 2-chlorotritylchloride resin (1 eq., 0.25 mmol) as described above. The remainder of the linear peptide was synthesized and acetylated as described above, replacing Fmoc-N-methyl-threonine-OH with Fmoc-Dap(Boc)-OH. The peptide was cleaved from the resin while removing the Boc protecting group by adding a solution of TFA in DCM (30% v/v containing 2.5% v/v triisopropylsilane; 10 mL total) and shaking at room temperature for 2 h. This solution was concentrated, washed and cyclized as described above to afford crude peptide.

#### Synthesis of 20

The linear peptide was synthesized as described above, replacing Fmoc-N-methyl-threonine-OH with Fmoc-Lys(Mmt)-OH. The peptide was acetylated as described above. The peptide was cleaved from the resin while removing the Mmt protecting group by adding a solution of TFA in DCM (1% v/v containing 2.5% triisopropylsilane; 10 mL total) and shaking at room temperature for 2 h. This solution was concentrated, washed and cyclized as described above to afford crude peptide.

#### Synthesis of 23, 24, and 25

Peptides were synthesized as described above, replacing Pro_2_ (**23**), Pro_5_ (**24**) or Pro_8_ (**25**) with Fmoc-homoproline-OH.

#### Synthesis of 26 and 27

Peptides were synthesized as described above, replacing Pro_8_ with either Fmoc-4-benzyloxy-proline (**26**) or Fmoc-4-*tert*-butyoxy-proline (**27**).

#### Synthesis of 28, 29, and 30

Peptides were synthesized as described above, replacing Pro_2_ (**28**), Pro_5_ (**29**), or Pro_8_ (**30**) with *trans*-*N*-Fmoc-4-propargyloxy-L-proline. The details for the synthesis of *trans*-*N*-Fmoc-4-propargyloxy-L-proline are described below, with further characterization information in the Supplementary Information.

#### Synthesis of 31 and 32

Peptides were synthesized as described above, replacing Me-Val_1_ with Fmoc-Pra-OH (**30**) or 4-pentynoic acid (**31**).

#### Synthesis of 33

The linear peptide was synthesized as described above, except Boc-valine-OH was used as the final residue in place of Fmoc-*N*-methyl-valine-OH. The peptide was esterified, cleaved, and cyclized as described above. The cyclization reaction mixture was concentrated and washed with hexanes (3 x 5 mL) followed by a 1:1 solution of hexanes in cold diethyl ether (10 mL), then the Boc protecting group was removed by adding a solution of TFA in DCM (30% v/v containing 2.5% v/v triisopropylsilane; 10 mL total) and shaking at room temperature for 2 h. The solution was concentrated and washed with hexanes (3 x 5 mL) followed by a 1:1 solution of hexanes in cold diethyl ether (10 mL) to afford crude peptide.

#### Synthesis of 34

The linear peptide was synthesized as described above, except Fmoc-Orn(Boc)-OH was appended after the *N*-terminal Me-Val_1_. The remainder of the peptide was synthesized as described for **33**.

#### Synthesis of 35

The linear peptide was synthesized as described above. After adding the final residue (Fmoc-methyl-valine), the Fmoc protecting group was removed and the resin was shaken with a solution of NBD-Cl (3 eq., 0.75 mmol) and DIEA (15 eq., 3.75 mmol) in DMF (7 mL) for 2 h). The peptide was then washed, esterified, cleaved, and cyclized as described above.

#### Synthesis of 36

The linear peptide was synthesized as described above, replacing Pro_2_, Pro_5_, and Pro_8_ with *trans*-*N*-Fmoc-4-propargyloxy-L-proline (the synthesis of which is described below), Fmoc-HomoPro-OH, and Fmoc-4-benzyloxy-Pro-OH respectively. Fmoc-Orn(Boc)-OH was appended after the *N*-terminal Me-Val_1_. The remainder of the peptide was synthesized as described for **33**.

#### Purification of peptides

All peptides were purified via reverse-phase high performance liquid chromatography (RP-HPLC) with a Waters 1525 Binary HPLC Pump and a Waters 2489 UV/Vis detector on either a Phenomenex Luna 10 µm C8(2) 100 Å 250 x 21.2 mm or a Phenomenex Aeris 5 µm PEPTIE XB-C18 100 Å 250 x 21.2 mm column using gradient elution of H2O/ACN + 0.1% TFA at 10 mL/min. Fractions containing the desired compound were concentrated using a rotary evaporated then lyophilized to dryness using a Labconco FreeZone lyophilizer. Peptide identities were confirmed via matrix-assisted laser desorption time-of-flight mass spectrometry (MALDI-MS) using a Shimadzu 8020. Peptides were analyzed for purity using the same RP-HPLC system and a Phenomenex Gemini 5 µm C18 110 Å 250 x 4.6 mm column. All peptides used in experiments had ≥ 95% purity as determined by integration of analytical HPLC chromatograms measured at an absorbance of 220 nm.

#### Synthesis of *trans*-*N*-Boc-4-propargyloxy-L-proline

A solution of N-Boc-trans-4-hydroxy-L-proline (1 g, 4.32 mmol) in anhydrous tetrahydrofuran (10 mL) under nitrogen was added via syringe to a 100 mL round bottom flask under nitrogen containing sodium hydride (NaH, 60% oil dispersion) (0.38 g, 9.51 mmol) in anhydrous THF (10 mL). The reaction mixture was cooled to 0 °C and stirred for 1 hr. Propargyl bromide (0.43 mL, 4.76 mmol) was added neat to the reaction mixture, and the reaction was stirred overnight at room temperature (RT). The reaction was quenched with H_2_O (5 mL) and acidified to pH 3-4 with 10% hydrochloric acid. The resulting mixture was extracted with ethyl acetate (EtOAc) (3 × 20 mL). The combined organic layers were washed with brine (2 × 20 mL), dried over anhydrous sodium sulfate (Na2SO4), filtered, and concentrated in vacuo. The crude material was purified by silica gel flash column chromatography using a mixture of dichloromethane (DCM), methanol (MeOH) and acetic acid (AcOH) (96:3:1). The desired fractions were collected, and the solvent was removed in vacuo to yield *trans*-*N*-Boc-4-propargyloxy-L-proline in 95% yield (1.1g).

^1^H NMR (400 MHz, CDCl3): δ 4.52 - 4.25 (m, 2H), 4.22 - 4.07 (m, 2H), 3.72 - 3.47 (m, 2H), 2.49 - 2.28 (m, 2H), 2.28 - 2.10 (m, 1H), 1.45/1.41* (s, 9H) (*indicates minor rotamer).

^13^C NMR (400 MHz, CDCl3): δ 178.2, 174.7, 156.4, 153.8, 81.8, 80.8, 79.2, 76.0, 75.7, 74.9, 57.8, 57.7, 56.5, 56.4, 51.8, 51.1, 36.5, 34.2, 28.3.

#### Synthesis of *trans*-*N*-Fmoc-4-propargyloxy-L-proline

To the prepared *trans*-*N*-Boc-4-propargyloxy-L-proline (1.1 g, 4.08 mmol) in a round-bottom flask equipped with a stir bar was added a solution of trifluoracetic acid (TFA) and DCM (20%:80%, 10 mL). The reaction mixture was stirred at RT for 3 hrs, the TFA/DCM was removed and dried *in vacuo*. The residue was dissolved in 10 mL of water, and the flask was allowed to cool to 0 °C. To the flask, NaHCO_3_ was added (1g, 12.84 mmol) and the mixture was stirred for 15 minutes, followed by addition of a solution of N-(9-Fluorenylmethoxycarbonyloxy) succinimide (FmocOsu) (1.5 g, 4.5 mmol) in acetone (10 mL). The reaction mixture was stirred overnight at RT, followed by acidification to pH 1-2 with 10% HCl. Subsequently, the reaction mixture was extracted with deionized H_2_O (25 mL) and EtOAc (3 × 20 mL). The combined organic layers were washed with brine (2 × 30 mL), dried over anhydrous Na_2_SO_4_, filtered, and concentrated *in vacuo*. The crude material was purified by silica gel flash column chromatography using a mixture of DCM:MeOH:AcOH (96:3:1) as eluent. The desired fractions were collected, and the solvent was removed *in vacuo* to yield *trans*-*N*-Fmoc-4-propargyloxy-L-proline in 85% yield as a white solid.

^1^H NMR (400 MHz, CDCl3): δ 7.74 (dd, J = 28.4, 7.5 Hz, 2H), 7.62 -7.49 (m, 2H), 7.46 - 7.20 (m, 5H), 4.54 - 4.23 (m, 5H), 4.20 - 4.08 (m, 2H), 3.81 - 3.59 (m, 2H), 2.54 - 2.12 (m, 3H)

^13^C NMR (400 MHz, CDCl3): δ 177.2, 175.2, 156.1, 154.5, 143.9, 143.7, 143.6, 141.3, 127.8, 127.6, 127.1, 125.0, 120.0, 119.9, 79.1, 75.5, 75.1, 75.0, 68.1, 67.7, 58.0, 57.3, 56.5, 51.7, 51.6, 47.2, 47.1, 36.7, 34.7.

### Computational Methods

Sequence alignments were generated using UniProt, TCoffee, and Espript. Percent similarities were calculated using Espript. Structure alignments and molecular visualization were performed using UCSF ChimeraX. The binding pocket of *M. tuberculosis* was defined as any residue containing an atom within 5 Å of **GMe** in the DnaN-**GMe** crystal structure (PDB: 5AGU). For **Fig. 1c – e**, the Pol III complex from *E. coli* consisting of DNA bound to the α, β, and ε subunits (PDB: 5FKW) was used as a reference to gauge potential **GMe**-pol III interactions as no pol III crystal structure from mycobacteria was available. **GMe** co-crystallized DnaN from *E. coli* (PDB: 8CIX) was aligned with this structure and the duplicate DnaN was hidden from view to aid visualization.

### Biological Methods

#### Mycobacteria cell culture

Cultures of *M. smegmatis* strain mc^2^155 (ATCC 700084) were grown using a 0.1% v/v inoculation from 50% glycerol stocks of stationary phase cells. Cells were grown in 7H9 media containing glycerol (0.5%) and tween-80 (0.05%) supplemented with 10% v/v albumin-dextrose-catalase (ADC) solution (sterilized H_2_O containing 50 mg/mL BSA, 20 mg/mL dextrose, and 0.03 mg/mL catalase). All cultures were grown in 15 mL culture tubes at 37 °C while shaking at 250 rpm.

#### MIC assays in *M. smegmatis*

Briefly, 3 mL of an *M. smegmatis* culture was prepared as described above and grown for 24 h to an OD_600_ of ∼0.5. Peptide stocks (25.6 mM in DMSO) were diluted in 7H9 media to 2X the highest concentration; all dilutions in this assay were done using ADC enriched 7H9 media. 200 µL of each stock was added in triplicate to a 1.2 mL deep well 96-well plate, after which twofold serial dilutions were performed. The cell culture was diluted 50-fold to an OD_600_ of ∼0.01, then 200 µL of this solution was added to each well to give a final volume of 400 μL. One row of the plate was reserved for wells without peptides (growth control) and wells without peptides or cells (sterility control). The plates were covered with a porous coverslip and incubated at 37 °C while shaking at 250 RPM for 40 h. 200 µL from each well was transferred to a clear flat-bottom 96 well plate. MIC values were determined as the lowest concentration of peptide that gave OD_600_ < 0.1 read using a Biotek Synergy H1 plate reader. MIC data are the average of three technical replicates which were shown to be equal between 3 biological replicates.

#### MIC assays in M. tuberculosis

The double auxotroph *M. tuberculosis* mc^2^6206 strain (H37Rv Δ*panCD* Δ*leuCD*1) was provided by Dr. William Jacobs and was cultured in Middlebrook 7H9 supplemented with 0.5% glycerol, 10% Middlebrook Oleic Albumin Dextrose Catalase (OADC; BD), and either 0.05% Tyloxapol, for strain maintenance, or 0.05% Tween-80, for experimental manipulation.^51,52^ Growth media were additionally supplemented with 50 μg/mL L-leucine and 50 µg/mL pantothenic acid. Bacterial stocks were prepared by directly freezing at -80 °C *M. tuberculosis* culture at OD_600_ ∼0.5. MIC assays were performed as described for *M. smegmatis* by serially diluting the compounds in 100 µL of supplemented media and adding 100 µL of *M. tuberculosis* culture diluted to OD_600_ ∼0.02 (final volume of 200 µL). Plates were incubated at 37 °C while shaking at 250 RPM for 10 days. OD_600_ was measured by fixing 150 µL of each well in 50 µL of a solution of 16% paraformaldehyde in PBS. MIC values were determined as the lowest concentration of each compound that resulted in no growth as determined visually and by OD_600_. MIC data shown are the average of three biological replicates.

#### MIC assays in *E. coli*

^53^ Briefly, an *E. coli* strain ATCC 25922 culture was grown for 16 h from a glycerol stock stab in 3 mL of Mueller-Hinton II broth (all dilutions in this assay were done using Mueller-Hinton II broth). A 3 mL subculture was made using a 1% inoculation from the overnight culture and grown for 6 h to an OD_600_ ∼1. This culture was diluted to an OD_600_ of ∼0.075, then further diluted 50-fold. 96-well plates containing peptides were prepared and serially diluted as described above, then 200 µL of diluted bacteria solution was added to each well. The plates were incubated for 16 h and MICs were measured as described above. For PMBN coincubation assays, PMBN was added to the specified wells at a final concentration of 10 µM. Assays were repeated in biological triplicates with one exemplary data set shown.

#### Lysate preparation

*M. smegmatis* cell lysate was prepared according to a previously established protocol.^54^ A 50 mL culture was grown to stationary phase (OD_600_ ∼1.6) as described above. The cell pellet was isolated and resuspended in 2 mL of a lysis buffer consisting of 50 mM Tris (pH 7.4), 200 mM NaCl, 0.5 mM CaCl_2_, 0.5 mM MgCl_2_, and 0.2% Triton X-100. The cell suspension was transferred to VWR® 2 mL pre-filled bead tubes (glass beads, 0.5 mm diameter) and cells were lysed by mechanical disruption using a VWR® 4-Place Mini Bead Mill Homogenizer (3 cycles of 1-min pulses at 5 m/s followed by 1 min on ice). Cell debris and beads were removed via centrifugation at 13,000 × g for 10 min at 4 °C. The supernatant was transferred to a fresh tube and further clarified by centrifugation at 15,700 × g for 5 min at 4 °C. The soluble lysate supernatant was further sterilized by filtration through a 0.2 µm PVDF membrane. The protein content of the lysate was quantified at 10.22 mg/mL using a ThermoFisher™ Pierce™ Dilution-Free™ Rapid Gold BCA Protein Assay kit.

#### Lysate and serum stability

Stability in the prepared *M. smegmatis* lysate was assessed for **GM**, **19**, **24** and **25**. The lysate was diluted to 2 mg/mL in PBS (pH 7.4). Compounds were incubated in 300 μL of lysate solution at a final concentration of 500 μM for 0, 15, or 30 min at 37 °C while shaking at 250 rpm. At each time point, 80 μL of the solution was aliquoted and diluted to 300 μL in a 50:50 solution of H_2_O/ACN containing 8.2% trichloroacetic acid (6.1 N) and 100 μM of an internal standard, Fmoc-L-phenylalanine. This mixture was incubated on ice for 5 min to precipitate proteins, followed by centrifugation at 21,000× g for 5 min. 250 μL of the supernatant was transferred to a fresh tube, which was again centrifuged at 21,000 × g for 5 min. 150 μL of the clarified supernatant was analyzed via RP-HPLC using the same column and system as described above. The area under the curve (AUC) for each compound was determined using Waters® Empower software. These values were normalized to the AUC of the Fmoc-L-phenylalanine standard from each run, and each compound was further normalized to its 0 min incubation AUC. Graphs were plotted using GraphPad Prism 8 software and curves were generated using one phase decay fittings. For serum stability experiments, the methods were repeated with incubations in 300 μL of human serum (Sigma-Aldrich®).

### Confocal imaging of *M. smegmatis*

A 3 mL *M. smegmatis* culture was prepared as described above and grown to stationary phase. Cells were harvested and centrifuged for 2 minutes at 4000 x g then washed three times in 3 mL PBST (PBS + 0.05% v/v Tween-20). After the final wash, the cells were resuspended in 3 mL PBST. NBD-GM was added to a final concentration of 10 µM and incubated for 30 minutes. 100 µL of cells was transferred to a round bottom 96-well plate, washed 3X with 100 µL PBST as described above, then resuspended in 100 µL of a solution of 4% formaldehyde in PBS. Cells were fixed for 15 minutes. 4 µL of fixed cells was transferred to a glass microscope slide and imaged on a Leica STELLARIS 8 Super resolution microscopy system (40x/1.3 oil-immersion lens) equipped with 405 nm, 488 nm, and 561 nm lasers. 488 nm was used as the excitation wavelength. We acknowledge the Keck Center for Cellular Imaging for the usage of the Leica STELLARIS 8 microscopy system. [PI: AP; NIH-OD030409].

#### Mammalian cell culture

J774A.1 macrophage cells were cultured in Dulbecco’s modified Eagle’s medium (DMEM) supplemented with 10% (v/v) FBS, 50 IU/mL penicillin, and 50 µg/mL streptomycin in a T75 flask and maintained in a humidified atmosphere of 5% CO_2_ at 37 °C.

### Confocal imaging of *M. smegmatis*-incubated macrophage cells

*M. smegmatis* cells were grown in liquid media as described above for approximately 24 hours to mid-log phase (OD_600_ ∼0.5 – 0.6). At this time, DADA was added to the culture to a final concentration of 100 μM, and the cells were grown for a further 16 hours at 37 °C while shaking at 250 rpm to stationary phase. The cells were harvested and centrifuged for 2 minutes at 4000 x g then washed three times in 3 mL PBST (PBS + 0.5% v/v Tween-20). J774A.1 cells were cultured as described above. On the day prior to the experiment, J774A.1 cells were seeded into 35 mm glass bottom microwell dishes and allowed to adhere. On the day of the experiment, J774A.1 cells were washed twice with 1X PBS. The PG-labeled *M. smegmatis* cells were resuspended in DMEM + 10% FBS containing no antibiotics and added to the washed J774A.1 cells at a multiplicity of infection of 100. The cell mixture was incubated at 37°C for 1 hour to induce phagocytosis and subsequently washed five times with 1X PBS to remove extracellular bacteria. The cells were then incubated with a solution containing 10 µM **35** in DMEM containing no FBS and no antibiotics for 30 min at 37°C. The cells were washed three times with 1X PBS and fixed for 30 min with 4% formaldehyde in 1X PBS. Cells were imaged as described above. 488 nm was used as the excitation wavelength for **35** with emission gated to 550 - 700 nm. 405 nm was used as the excitation wavelength for DADA with emission gated from 420 - 480 nm.

## ANCILLARY INFORMATION

### SUPPORTING INFORMATION

Additional figures, synthetic schemes, a list of reagents, analytical HPLC chromatograms, and MALDI-MS spectra are included in the supporting information file (PDF). Molecular formula strings (CSV).

## ACKNOWLEDGEMENT

This study was supported by: the NIH grant 1R01AI178975-01 (M.M.P.), R35GM124893 (M.M.P.), and R01AI179080-01 (M.M.P. & M.S.S.); Gates Foundation (INV-080847 to M.S.S. and M.M.P.).

## AUTHOR CONTRIBUTIONS

A.S. and S.M.W. synthesized and characterized all peptides except **26** and **27** (Z.L.). A.S., R.D., S.M.W., I.L., and Y.C.L. carried out MIC assays. A.S., S.E.N., and S.B. carried out microscopy experiments. A.S. and S.M.W. carried out serum and lysate stability experiments. V.C. synthesized *trans*-*N*-Fmoc-4-propargyloxy-L-proline. A.S. prepared figures and tables. A.S. and M.M.P. wrote the manuscript. J.C., M.S.S. and M.M.P. directed the research. All authors have given approval to the final version of the manuscript.

## NOTES

The authors declare no competing financial interest.

## ABBREVIATIONS

ACN: acetonitrile
Boc: *tert*-butoxycarbonyl
BSA: bovine serum albumin
DCM: dichloromethane
DIC: diisopropylcarbodiimide
DIEA: diisopropylethylamine
DMF: dimethylformamide
DMSO: dimethyl sulfoxide
Fmoc: 9-fluorenylmethoxycarbonyl
MDR: multidrug resistance
PBS: phosphate buffered saline
TFA: trifluoroacetic acid
THF: tetrahydrofuran.

