## Supporting Information for "Molecular Determinants Governing the Antitubercular Activity of Griselimycin"

### TABLE OF CONTENTS


| | |
|---|---|
| **SUPPLEMENTARY FIGURES** | S3 – S6 |
| **Figure S1.** Mycobacterial cell lysate and human serum stability assays with **GM** and **19** | S3 |
| **Figure S2.** Multiple sequence alignment of DnaN from *M. tuberculosis*, *M. smegmatis*, *E. coli*, and *H. pylori* | S4 |
| **Figure S3.** Mycobacterial cell lysate and human serum stability assays with **GM**, **24**, and **25** | S5 |
| **Figure S4.** *E. coli* minimum inhibitory concentration assay with **GM**, **33**, and **34** ± PMBN | S6 |
| **Figure S5.** Structure of the mycobacterial peptidoglycan label DADA | S7 |
| **Scheme S1.** General synthesis of griselimycin derivatives | S8 |
| **Scheme S2.** Synthesis of *trans*-*N*-Fmoc-4-propargyloxy-L-proline | S9 |
| **Materials** | S10 – S11 |
| **Characterization of Peptides** | S12 – S34 |
| **Characterization of *trans*-*N*-Boc-4-propargyloxy-L-proline** | S35 – S36 |
| **Characterization of *trans*-*N*-Fmoc-4-propargyloxy-L-proline** | S37 – S38 |
| **References** | S39 |

### SUPPLEMENTARY FIGURES

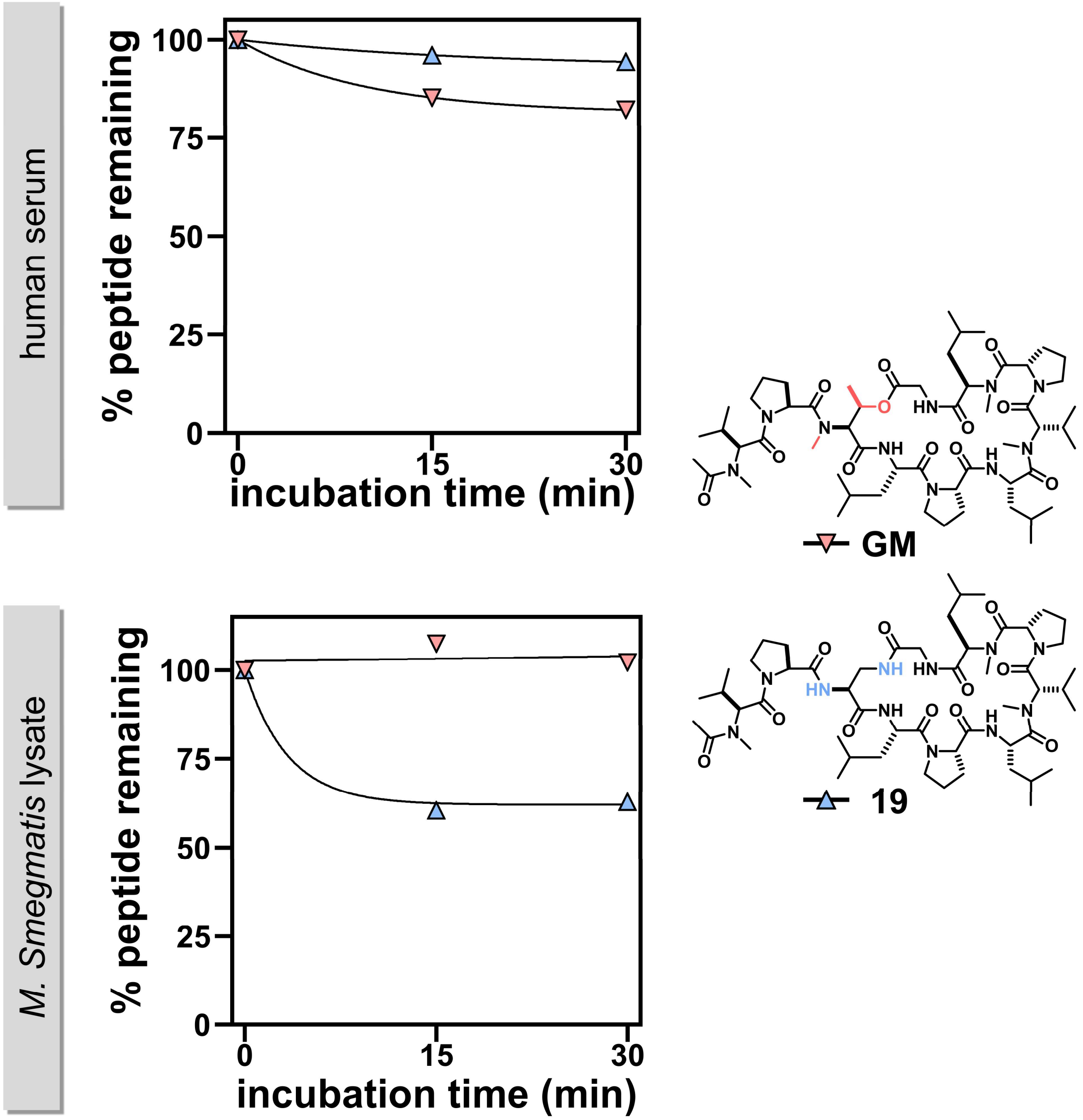


**Figure S1**. Time-course degradation of **GM** and **19** in human serum and *M. smegmatis* lysate. Peptides were incubated at 500 μM in either serum or lysate (diluted to a final protein concentration of 2 mg/mL) for 0, 15, or 30 minutes. Data points represent area under the curve of representative **GM** or **19** peaks from analytical HPLC chromatograms, normalized to the 0-minute incubation signal for each peptide. Each data set was fit to a one-phase decay curve using GraphPad Prism 8 software.

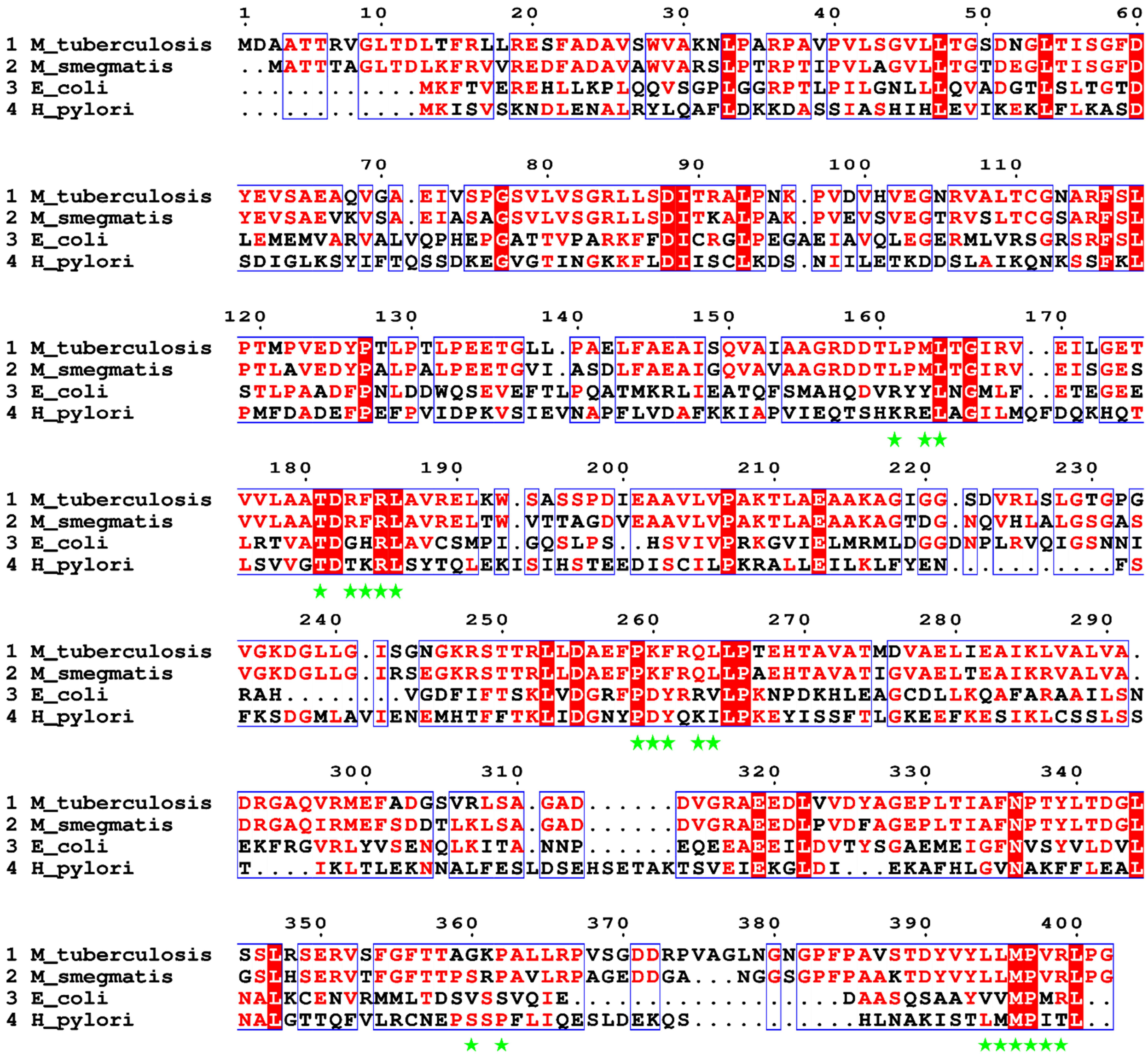


**Figure S2**. Multiple sequence alignment of DnaN from *M. tuberculosis*, *M. smegmatis*, *E. coli*, and *H. pylori* (PDB: 5AGU, 5AH2, 8CIX, and 5G48 respectively). Residues conserved between 2 – 3 organisms are in red text with continuous regions framed in blue. Strictly conserved residues are highlighted in red with white text. GM binding pocket residues in *M. tuberculosis* are marked with green stars.

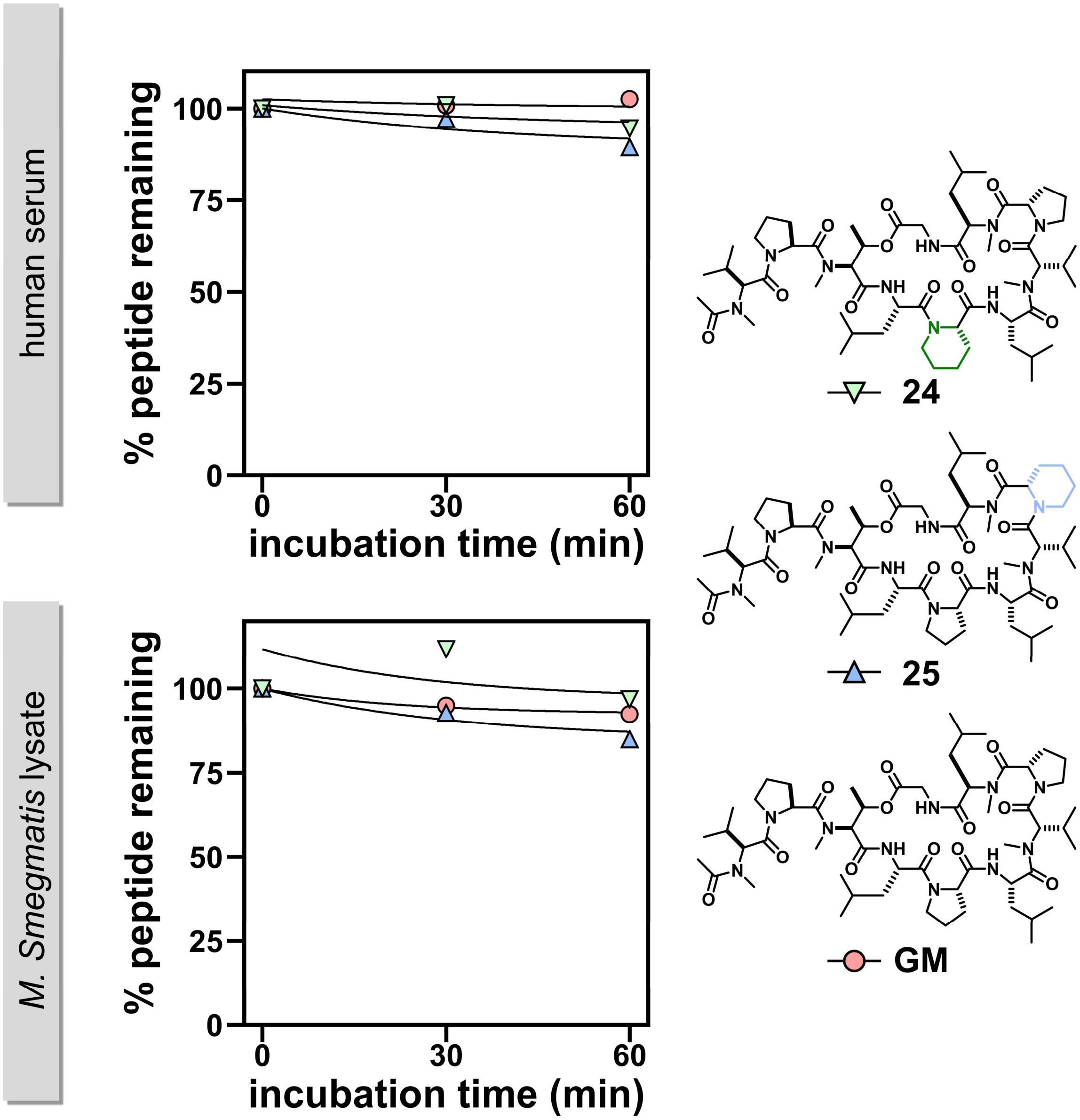


**Figure S3**. Time-course degradation of **GM**, **24**, and **25** in human serum and *M. smegmatis* lysate. Peptides were incubated at 500 µM in either serum or lysate (diluted to a final protein concentration of 2 mg/mL) for 0, 30, or 60 minutes. Data points represent area under the curve of representative **GM**, **24**, or **25** peaks from analytical HPLC chromatograms, normalized to the 0-minute incubation signal for each peptide. Each data set was fit to a one-phase decay curve using GraphPad Prism 8 software.

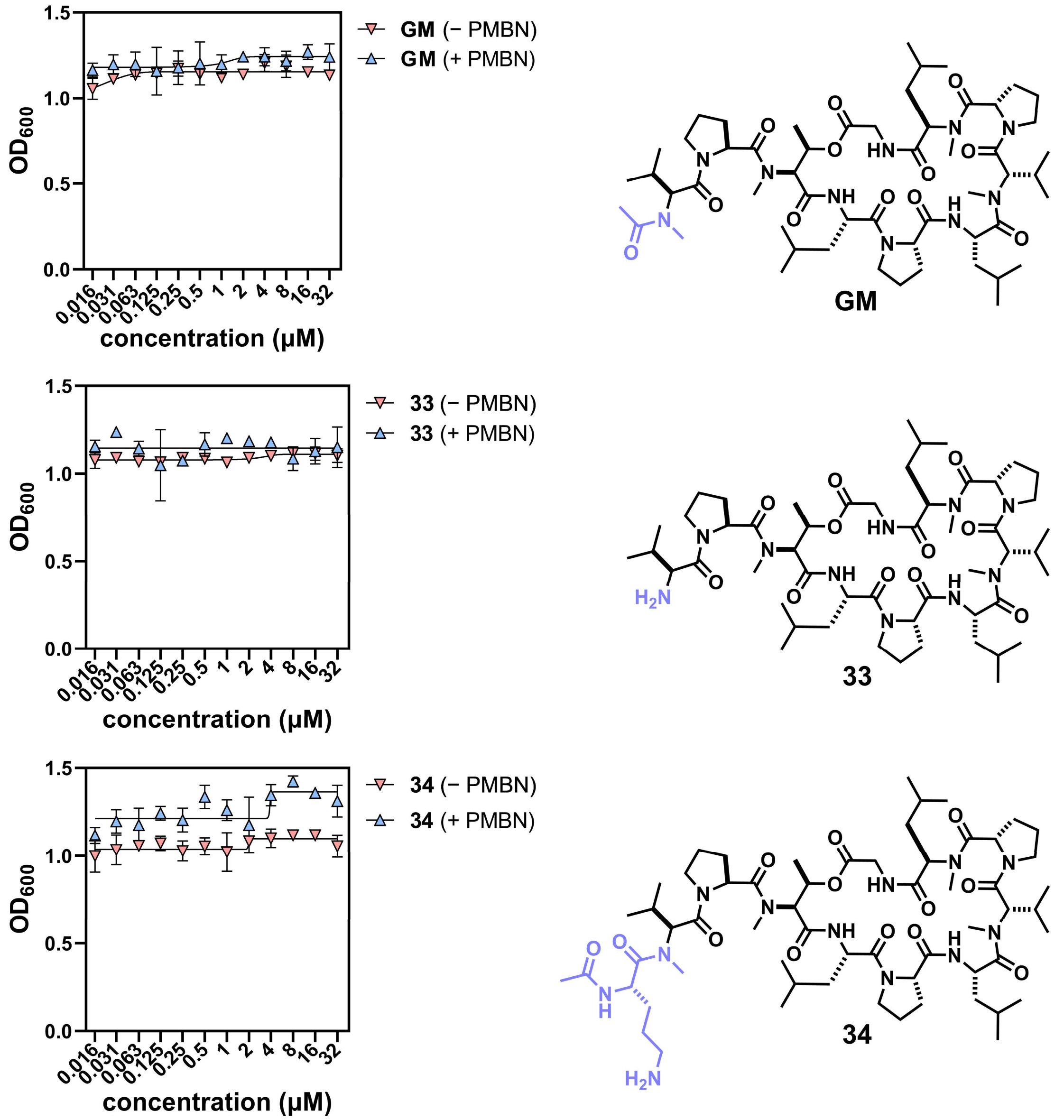


**Figure S4**. MIC assay of *E. coli* strain 25922 in the presence of varying concentrations of **GM**, **33**, or **34** with or without 10 μM PMBN coincubation. Data points represent the average of three replicates fit to a Boltzmann sigmoidal curve using GraphPad Prism 8 software; error bars represent ± one standard deviation.

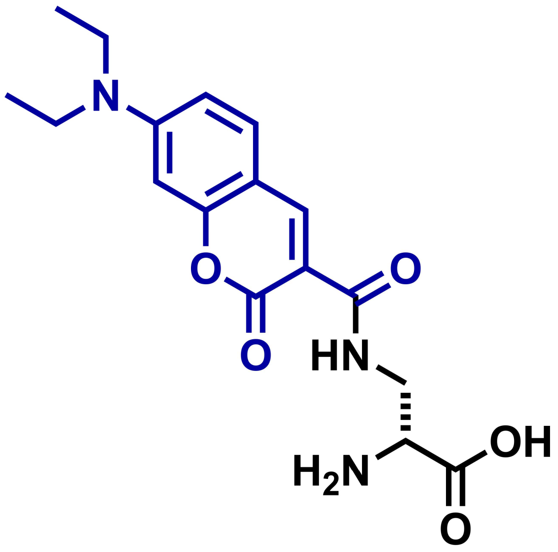


**Figure S5**. Structure of the mycobacterial peptidoglycan label DADA.[1]

**Scheme S1**. General scheme for griselimycin library synthesis

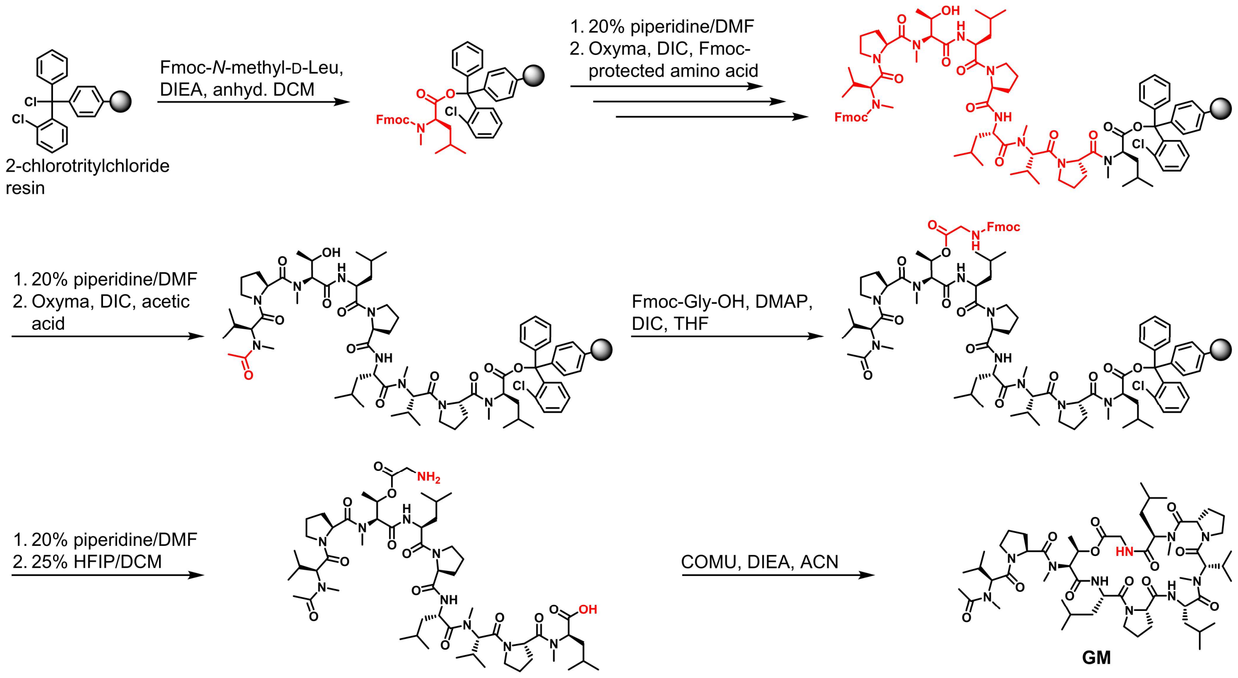

**Scheme S2.** Synthesis of *trans*-*N*-Fmoc-4-propargyloxy-L-proline

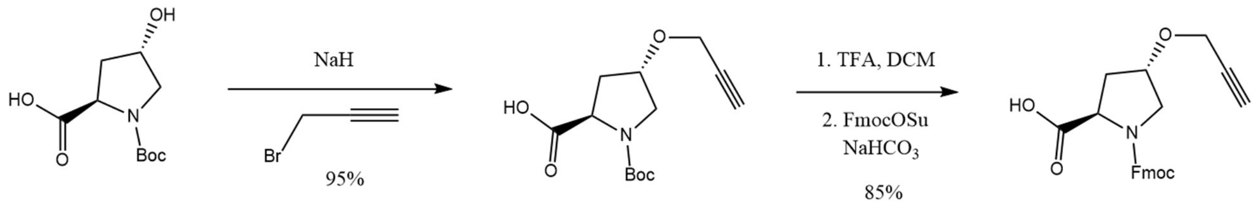

### Materials

| REAGENTS/MATERIALS | VENDOR SOURCE | CATALOG # |
|---|---|---|
| **Reagents/materials for cell culture and biological assays** | | |
| Middlebrook 7H9 media | VWR | 90003-876 |
| Catalase from bovine liver | Millipore Sigma | C1345-1G |
| D(+)-Glucose, anhydrous (dextrose) | Chem Impex | 00805 |
| Bovine serum albumin fraction V | Millipore Sigma | 10735078001 |
| Glycerol | Millipore Sigma | G7893 |
| Tween 80 | VWR | 97061-674 |
| Mueller-Hinton Broth | Millipore Sigma | 70192 |
| Rifampicin | Millipore Sigma | R3501 |
| Gentamycin | Millipore Sigma | 345814 |
| Griselimycin (GM) | Cayman Chemical | 39306 |
| PBS | ThermoFisher | 21600010 |
| Tween-20 | Millipore Sigma | 655204 |
| Pierce™ Dilution-Free™ Rapid Gold BCA Protein Assay | ThermoFisher Scientific | A55860 |
| Human Serum | Millipore Sigma | H4522 |
| VWR® Pre-Filled Bead Tubes, 2 ml, glass beads, 0.5 mm | VWR | 10158-606 |
| Polymyxin B nonapeptide hydrochloride | Millipore Sigma | P2076 |
| **Reagents for the synthesis and characterization of griselimycins** | | |
| *N*-Fmoc-amino acids | Chem Impex | Various |
| Fmoc-*N*-methyl-D-leucine-OH | Ambeed | A273782 |
| Fmoc-*N*-methyl-leucine-OH | Ambeed | A171951 |
| Fmoc-sarcosine-OH | Ambeed | A141362 |
| Fmoc-*N*-methyl-threonine-OH | Ambeed | A154318 |
| Fmoc-*N*-methyl-valine-OH | Ambeed | A278601 |
| Fmoc-phenylalanine(3,5-difluoro)-OH | Aapptec | UFF144 |
| Fmoc-Dap(Boc)-OH | Ambeed | 162558-25-0 |
| Boc-valine-OH | Ambeed | 13734-41-3 |
| Fmoc-HoPro-OH | Ambeed | A139951 |
| Fmoc-Orn(Boc)-OH | Ambeed | A120784 |
| Fmoc-Hyp(Bzl)-OH | Ambeed | A201678 |
| Fmoc-Hyp(tBu)-OH | Ambeed | A265743 |

| | | |
|---|---|---|
| NBD-Cl | Millipore Sigma | 163260 |
| *N*, *N′*-Diisopropylethylamine (DIEA) | Chem Impex | 00141 |
| *N*, *N′*-Diisopropylcarbodiimide (DIC) | Chem Impex | 00110 |
| Ethyl Cyano(hydroxyimino)acetate (Oxyma) | TCI Chemicals | E0847 |
| COMU | Millipore Sigma | 712191 |
| *N, N*-Dimethylformamide, ACS-grade and anhydrous | Millipore Sigma | 319937, 5.89565 |
| Dichloromethane ACS-grade and anhydrous | Millipore Sigma | D65100, 270997 |
| Tetrahydrofuran, anhydrous | Millipore Sigma | 186562 |
| Acetonitrile (ACN) ACS-grade and HPLC-grade | Millipore Sigma | 360457, 34851 |
| Piperidine | Millipore Sigma | 104094 |
| Trifluoroacetic acid, HPLC grade | Millipore Sigma | 302031 |
| Trifluoroacetic acid, reagent-grade | Chem Impex | 00289 |
| Hexafluoroisopropanol | Chem Impex | 00080 |

### Characterization of Peptides

The chemical structure, analytical HPLC chromatogram and MALDI-MS spectrum are shown for each synthesized peptide:

### GM

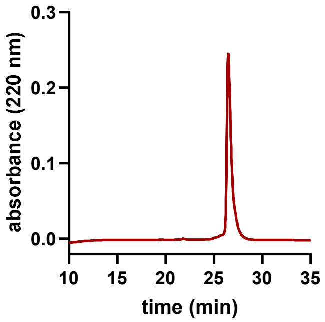


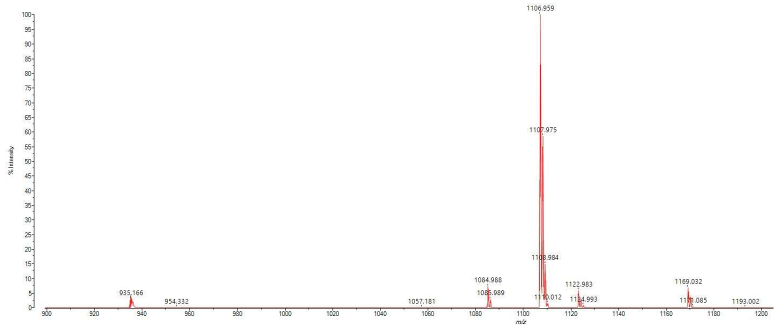


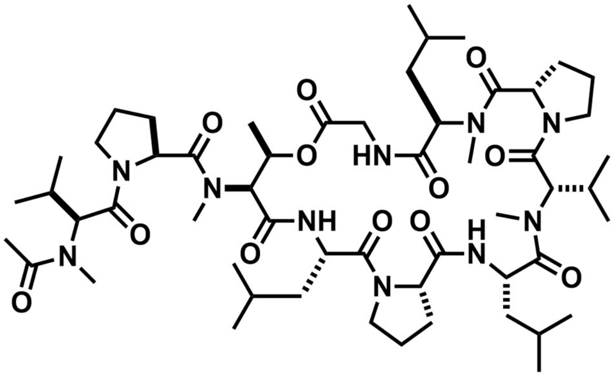

Chemical formula: $C_{55}H_{92}N_{10}O_{12}$. Exact mass (Da): 1084.690. m/z calculated for $[M + H]^{+}$: 1085.697; found: 1085.99.

**1**

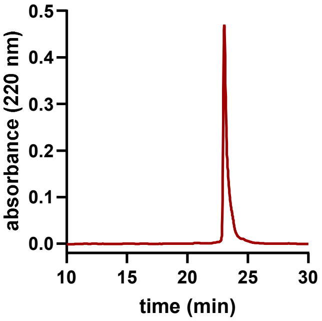


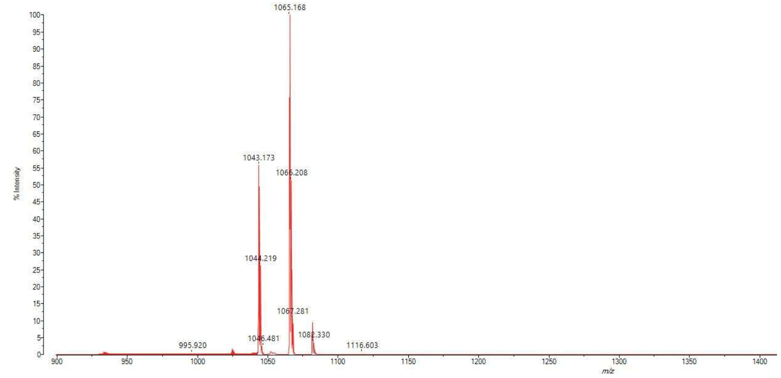


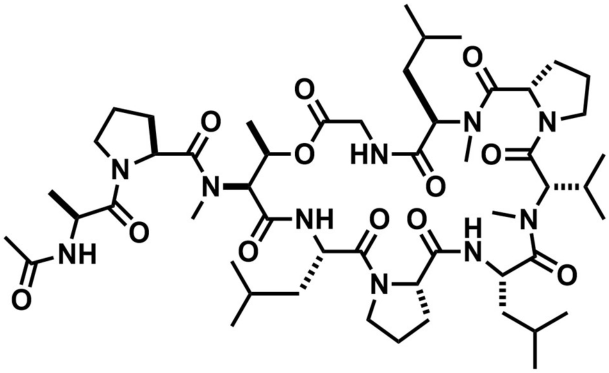

Chemical formula: $C_{52}H_{86}N_{10}O_{12}$. Exact mass (Da): 1042.643. m/z calculated for $[M + H]^+$: 1043.650; found: 1043.17.

**2**

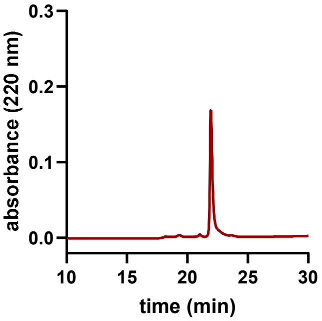


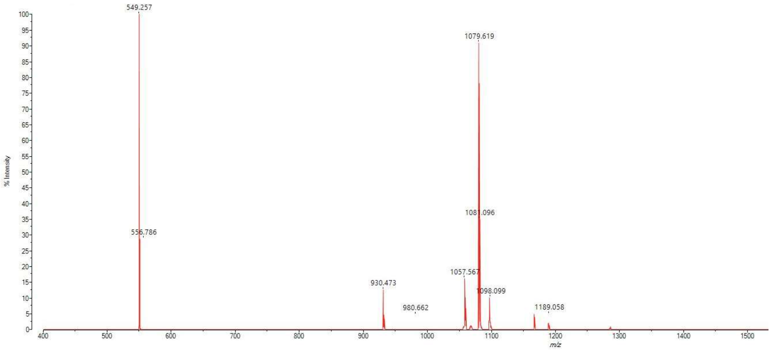


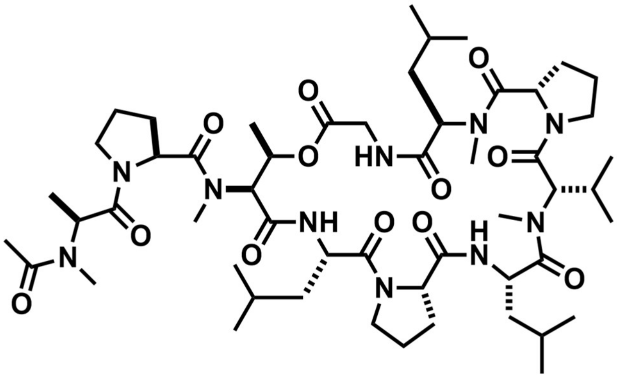

Chemical formula: $C_{53}H_{88}N_{10}O_{12}$. Exact mass (Da): 1056.658. m/z calculated for $[M + H]^+$: 1057.665; found: 1057.567.

**3**

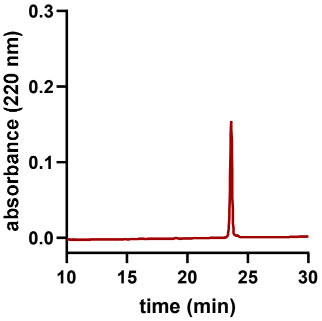


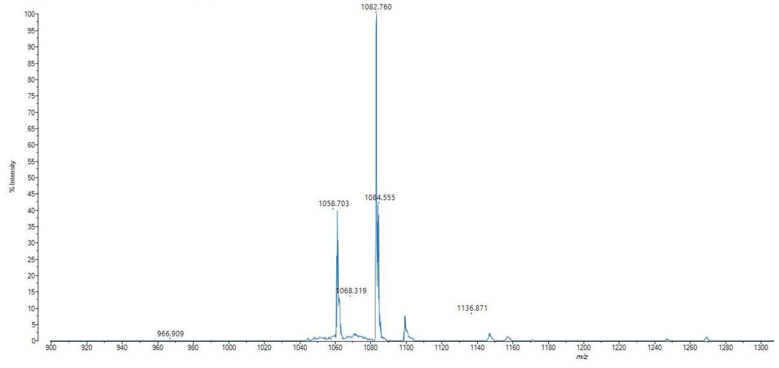


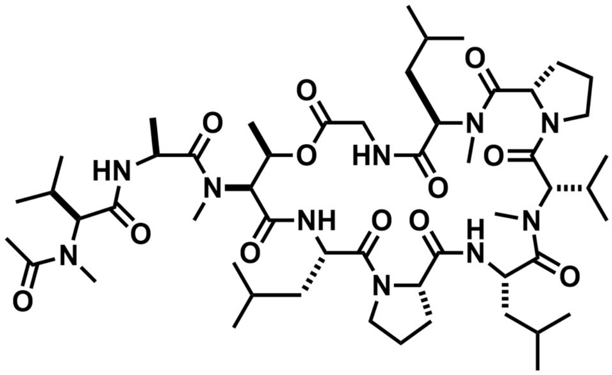

Chemical formula: $C_{53}H_{90}N_{10}O_{12}$. Exact mass (Da): 1058.674. m/z calculated for $[M + H]^+$: 1059.681; found: 1060.23.

**4**

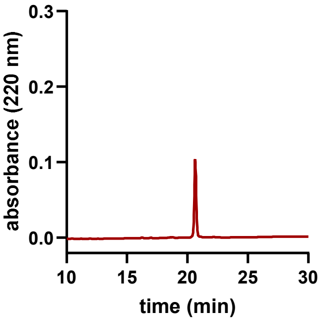


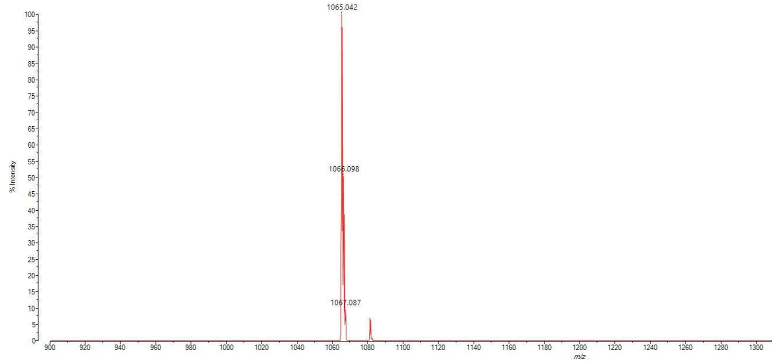

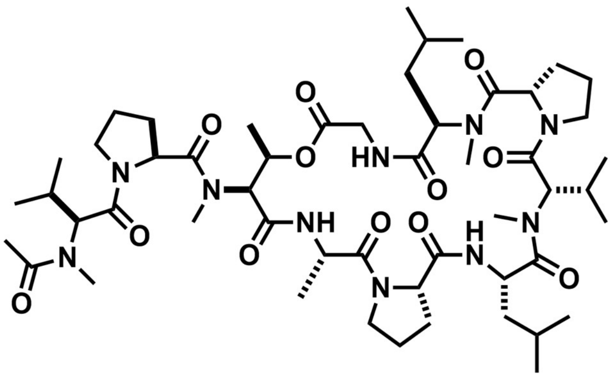

Chemical formula: $C_{52}H_{86}N_{10}O_{12}$. Exact mass (Da): 1042.643. m/z calculated for $[M + Na]^+$: 1065.633; found: 1065.04.

**5**

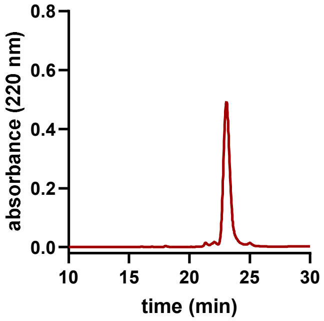


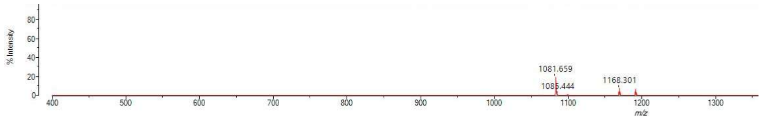


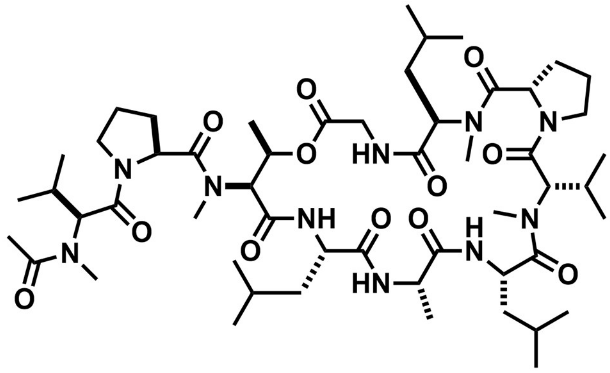

Chemical formula: $C_{53}H_{90}N_{10}O_{12}$. Exact mass (Da): 1058.674. m/z calculated for $[M + Na]^+$: 1081.664; found: 1082.02.

**6**

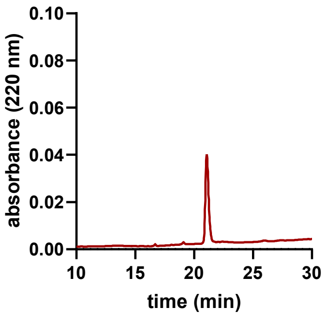


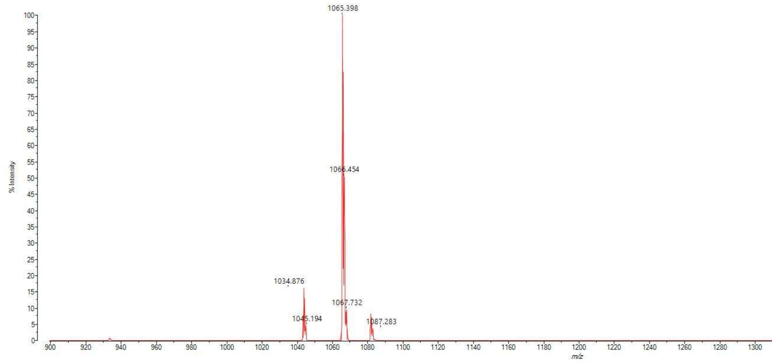


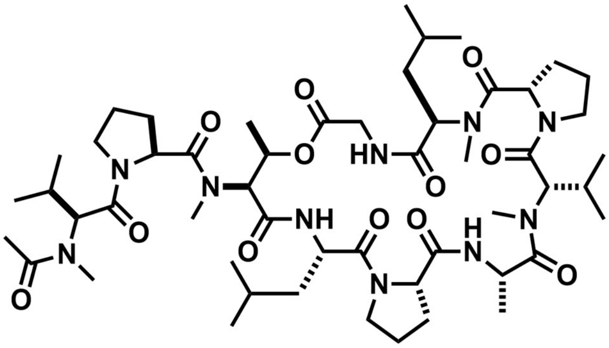


Chemical formula: $C_{52}H_{86}N_{10}O_{12}$. Exact mass (Da): 1042.643. m/z calculated for [M + Na]: 1065.633; found: 1065.40.

**7**

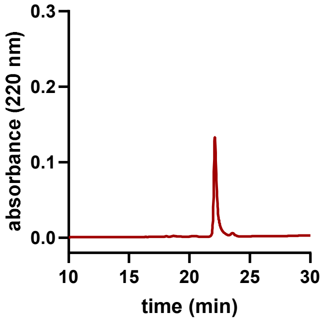


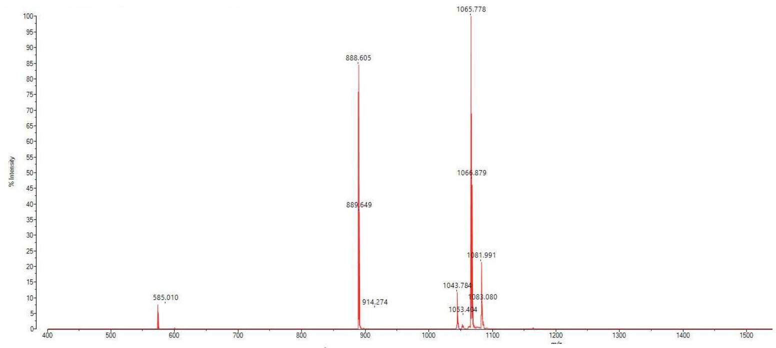


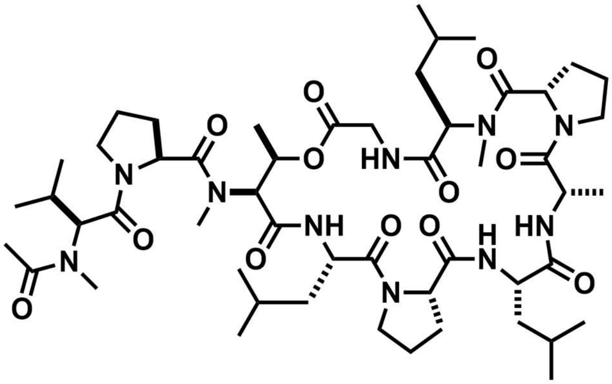

Chemical formula: $C_{52}H_{86}N_{10}O_{12}$. Exact mass (Da): 1042.643. m/z calculated for $[M + H]^+$: 1043.650; found: 1043.78.

**8**

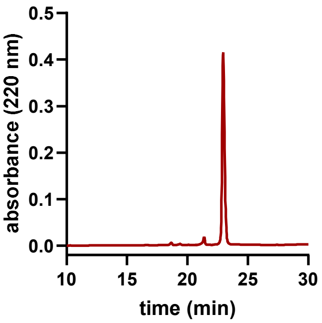


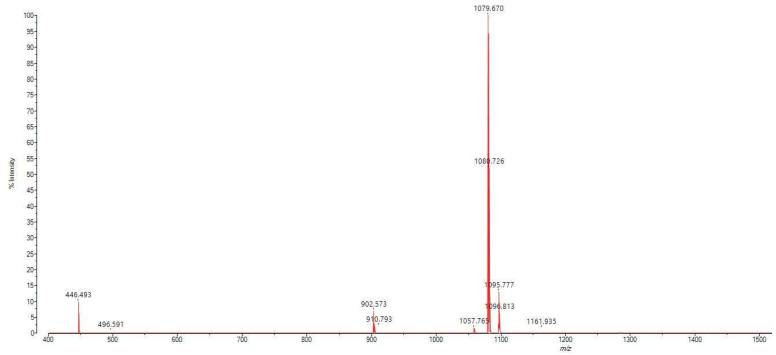


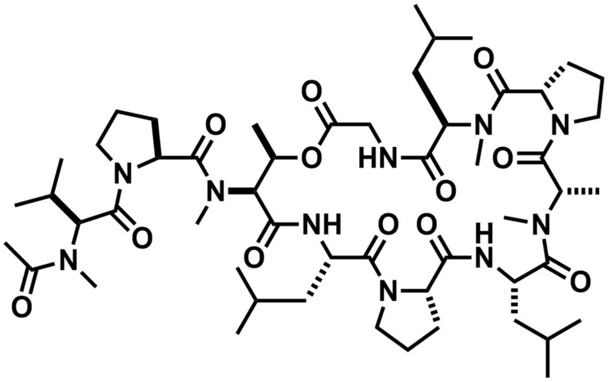


Chemical formula: $C_{53}H_{88}N_{10}O_{12}$. Exact mass (Da): 1056.658. m/z calculated for $[M + H]^+$: 1057.665; found: 1057.765.

**9**

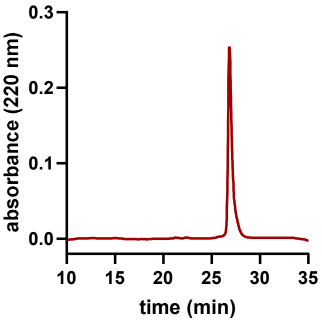


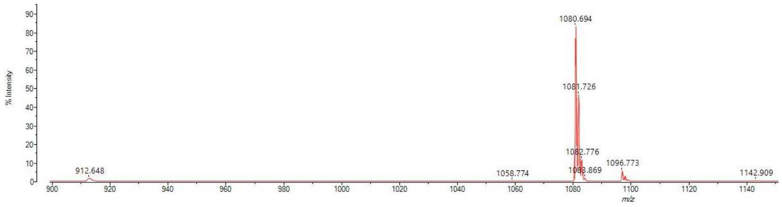

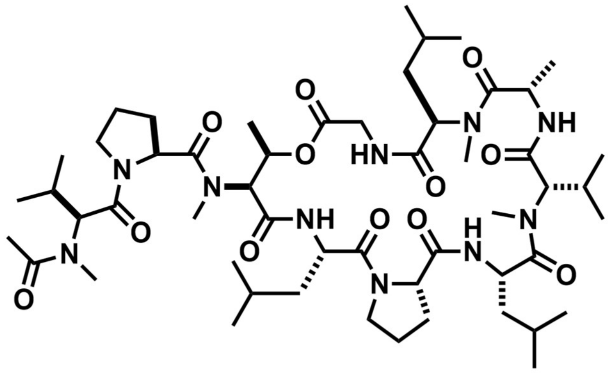

Chemical formula: $C_{53}H_{90}N_{10}O_{12}$. Exact mass (Da): 1058.674. m/z calculated for $[M + Na]^+$: 1081.664; found: 1081.726.

**10**

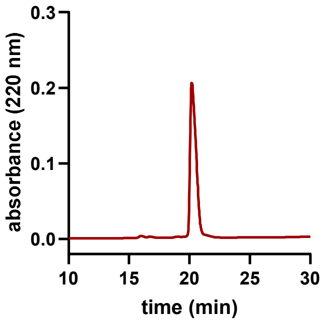


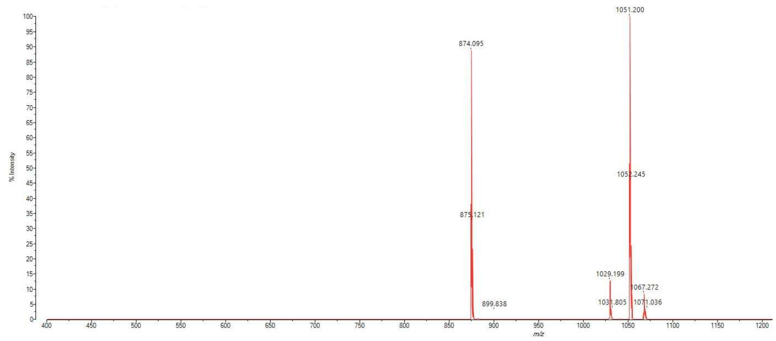


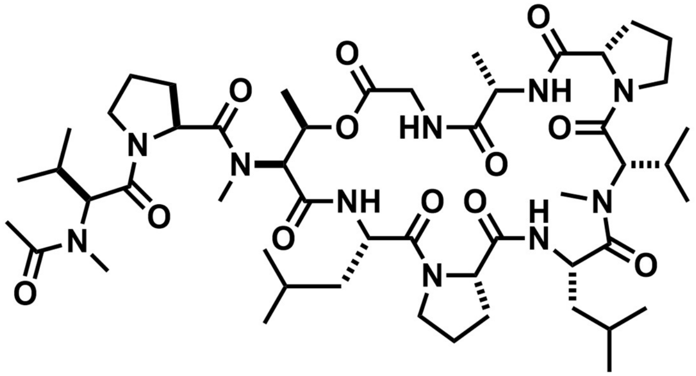

Chemical formula: $C_{51}H_{84}N_{10}O_{12}$. Exact mass (Da): 1028.627. m/z calculated for $[M + H]^+$: 1029.634; found: 1029.199.

**11**

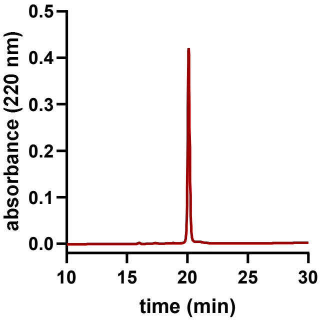


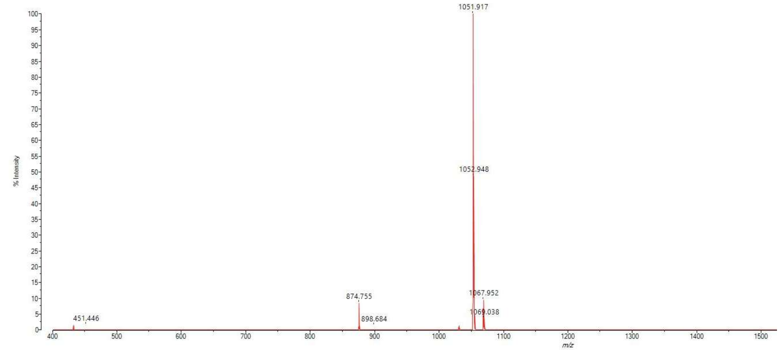


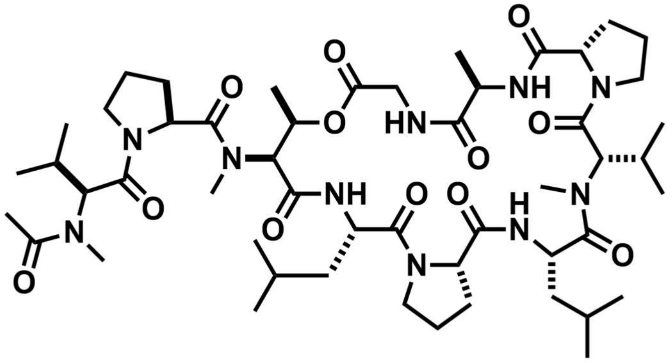

Chemical formula: $C_{51}H_{84}N_{10}O_{12}$. Exact mass (Da): 1028.627. m/z calculated for $[M + Na]^+$: 1051.617; found: 1051.917.

**12**

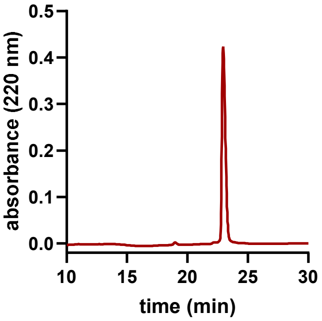


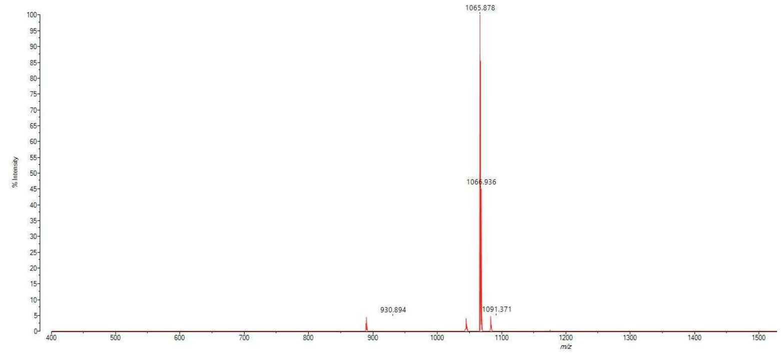


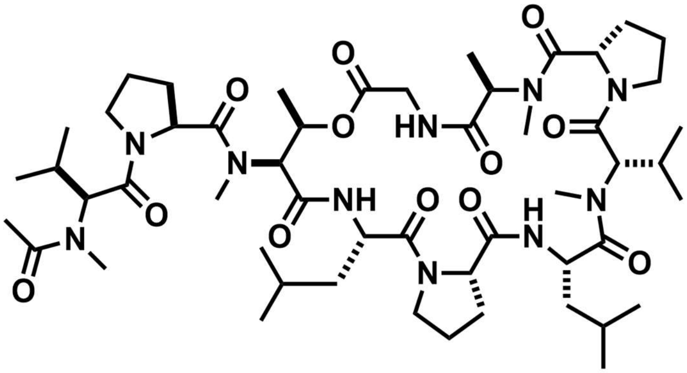

Chemical formula: $C_{52}H_{86}N_{10}O_{12}$. Exact mass (Da): 1042.643. m/z calculated for $[M + Na]^+$: 1065.633; found: 1065.878.

**13**

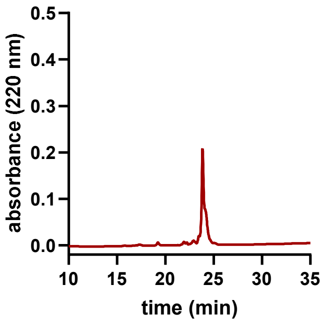


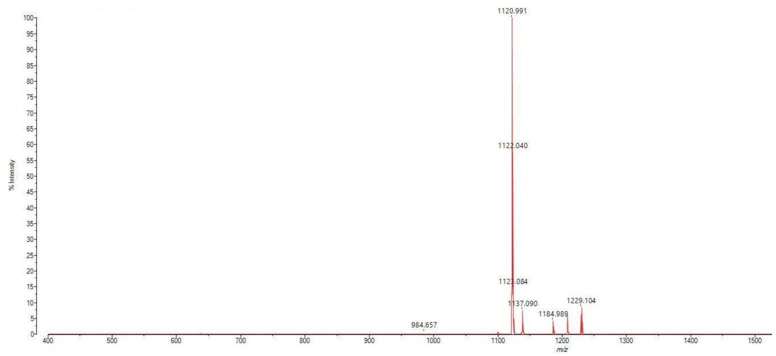


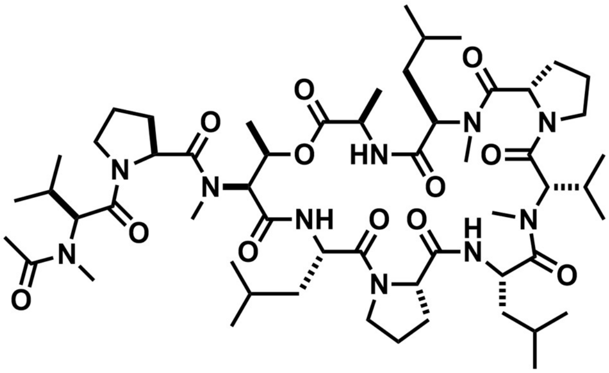

Chemical formula: $C_{56}H_{96}N_{10}O_{12}$. Exact mass (Da): 1098.705. m/z calculated for $[M + Na]^+$: 1121.695; found: 1120.991.

**14**

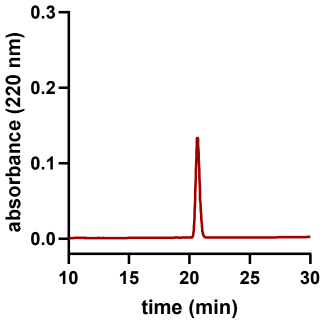


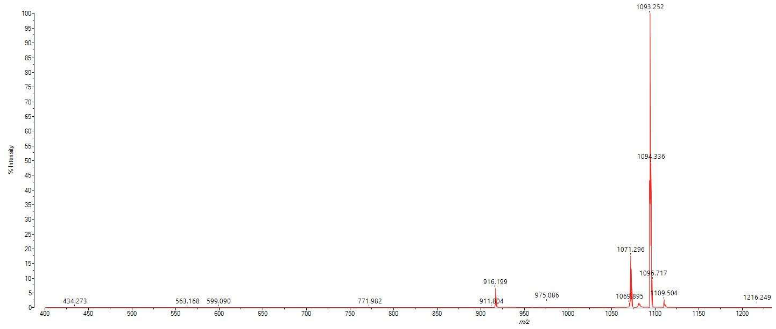

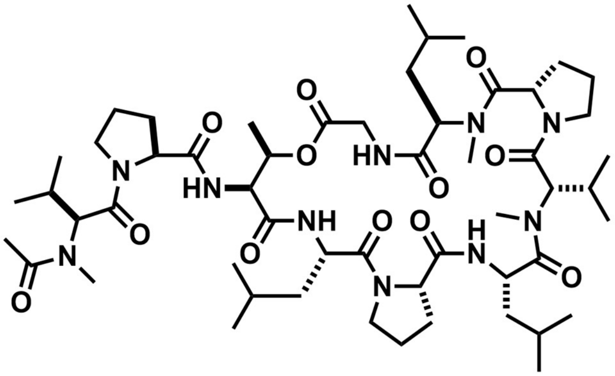

Chemical formula: $C_{54}H_{90}N_{10}O_{12}$. Exact mass (Da): 1070.674. m/z calculated for $[M + H]^+$: 1071.681; found: 1071.296.

**15**

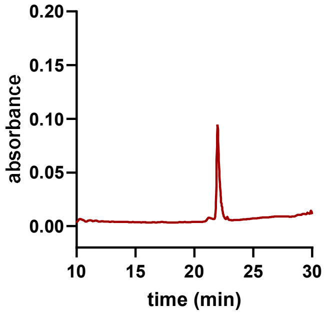


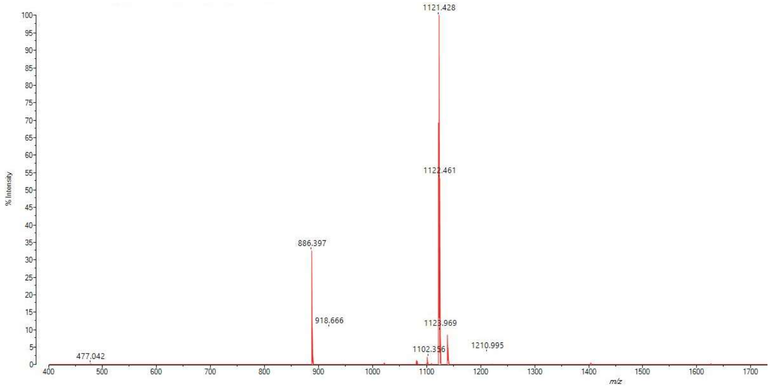


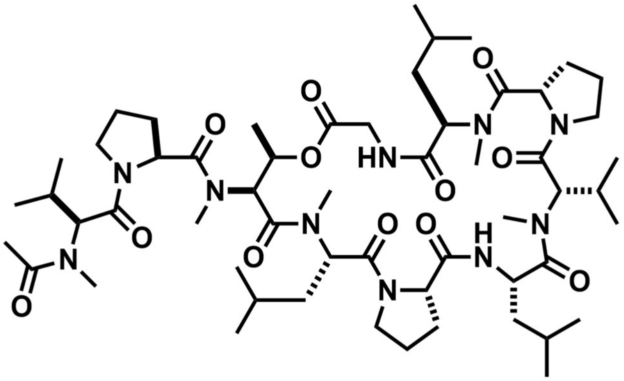

Chemical formula: $C_{56}H_{94}N_{10}O_{12}$. Exact mass (Da): 1098.705. m/z calculated for $[M + Na]^+$: 1121.695; found: 1121.428.

**16**

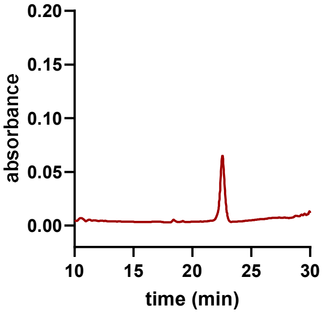


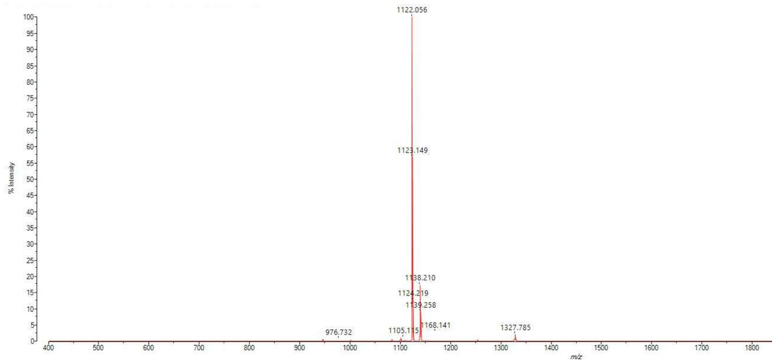


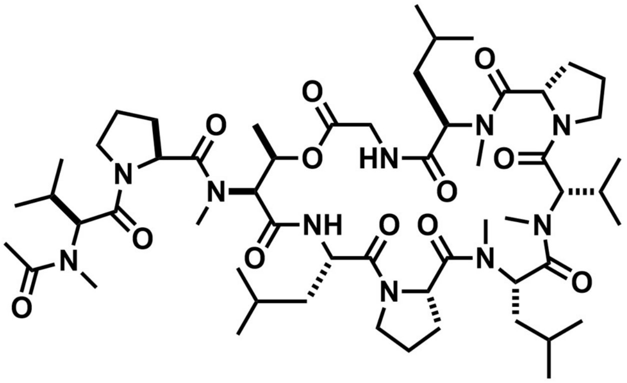

Chemical formula: $C_{56}H_{94}N_{10}O_{12}$. Exact mass (Da): 1098.705. m/z calculated for $[M + Na]^{+}$: 1121.695; found: 1122.056.

**17**

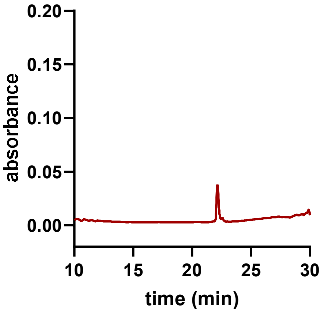


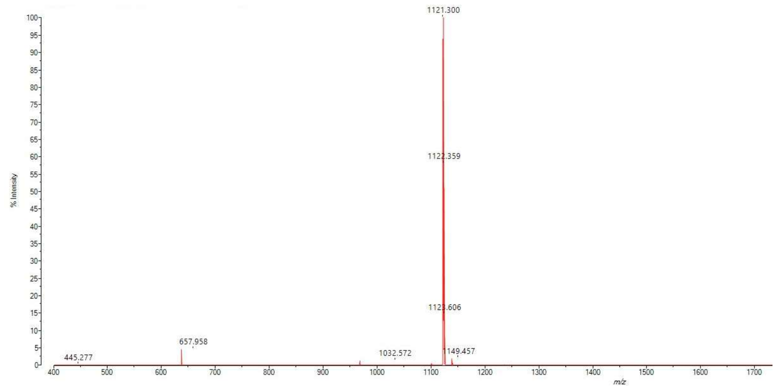


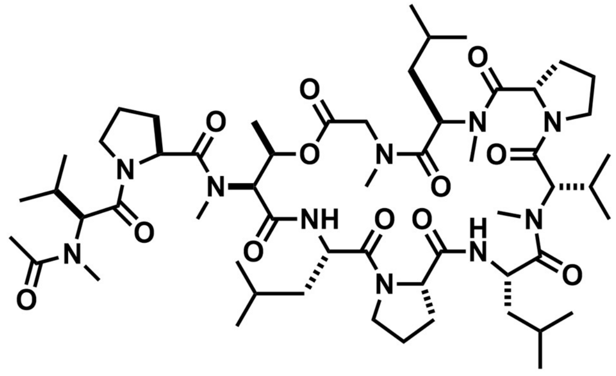

Chemical formula: $C_{56}H_{94}N_{10}O_{12}$. Exact mass (Da): 1098.705. m/z calculated for $[M + Na]^+$: 1121.695; found: 1121.300.

**18**

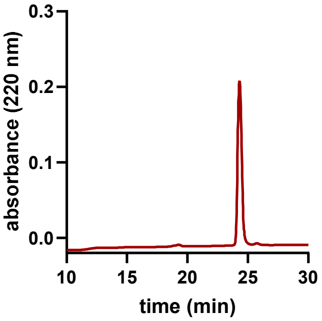


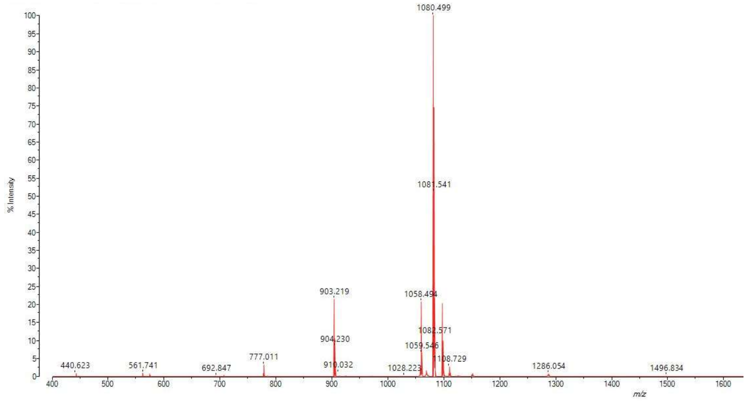


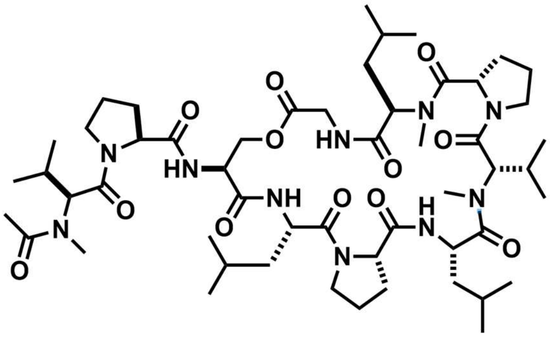

Chemical formula: $C_{53}H_{88}N_{10}O_{12}$. Exact mass (Da): 1056.658. m/z calculated for $[M + H]^+$: 1057.665; found: 1058.494.

**19**

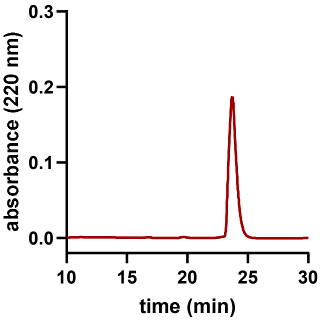


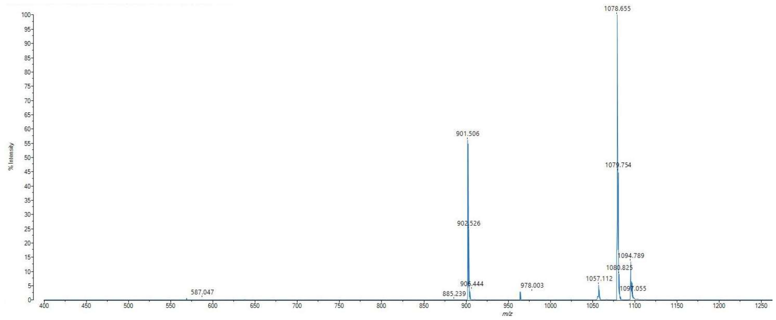

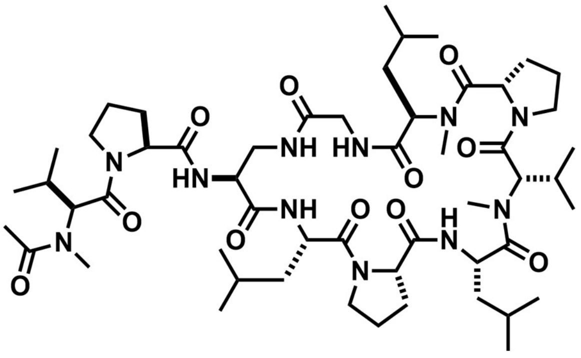

Chemical formula: $C_{53}H_{89}N_{11}O_{11}$. Exact mass (Da): 1055.674. m/z calculated for $[M + Na]^+$: 1078.664; found: 1078.655.

**20**

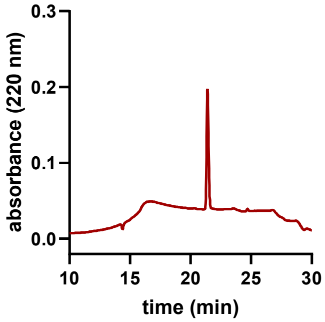


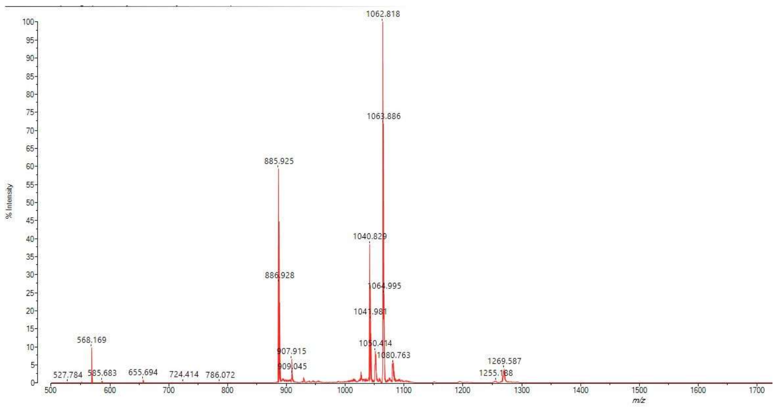


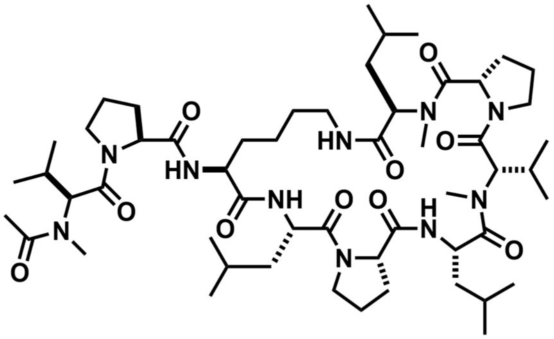

Chemical formula: $C_{54}H_{92}N_{10}O_{10}$. Exact mass (Da): 1040.700. m/z calculated for $[M + H]^+$: 1041.707; found: 1040.829.

**21**

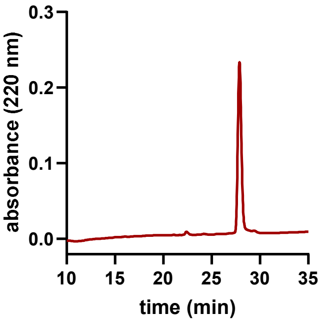


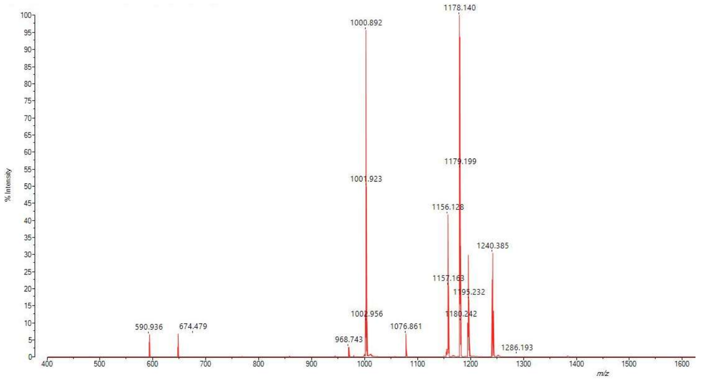


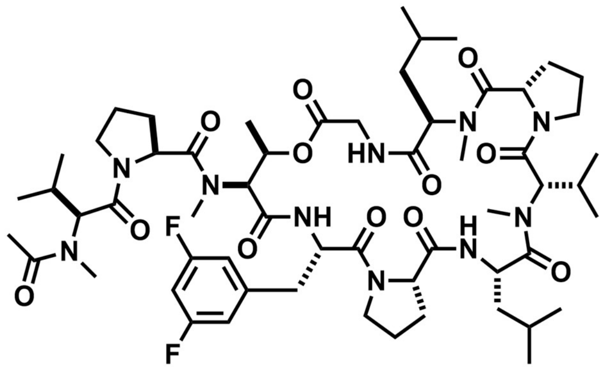

Chemical formula: $C_{58}H_{88}F_2N_{10}O_{12}$. Exact mass (Da): 1154.655. m/z calculated for $[M + H]^+$: 1155.662; found: 1156.128.

**22**

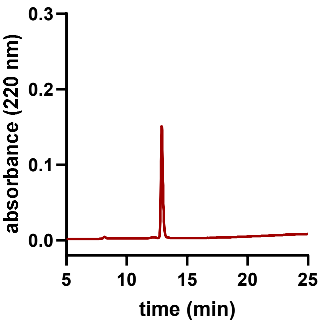


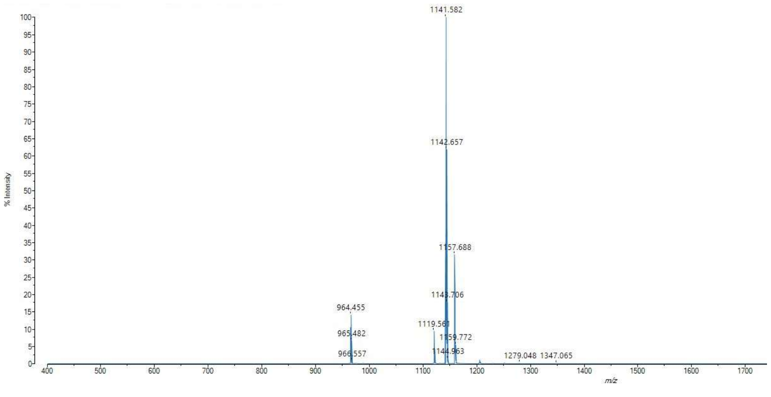

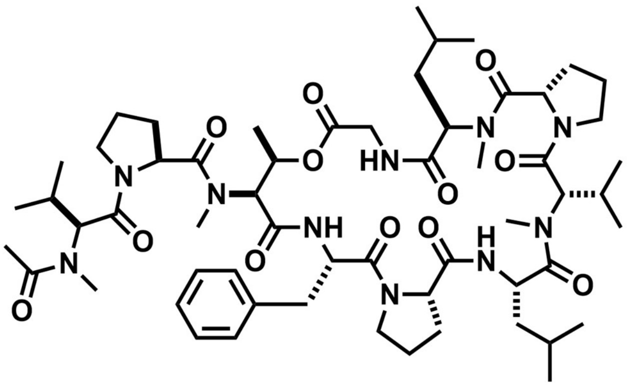

Chemical formula: $C_{58}H_{90}N_{10}O_{12}$. Exact mass (Da): 1118.674. m/z calculated for $[M + H]^+$: 1119.681; found: 1119.561.

**23**

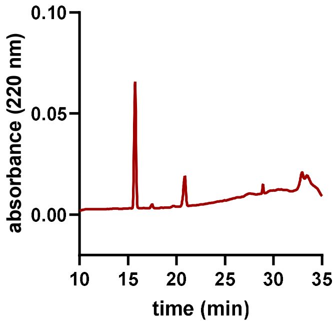


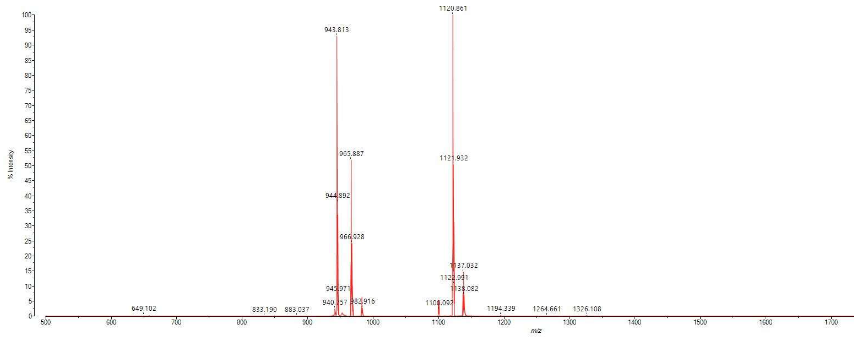

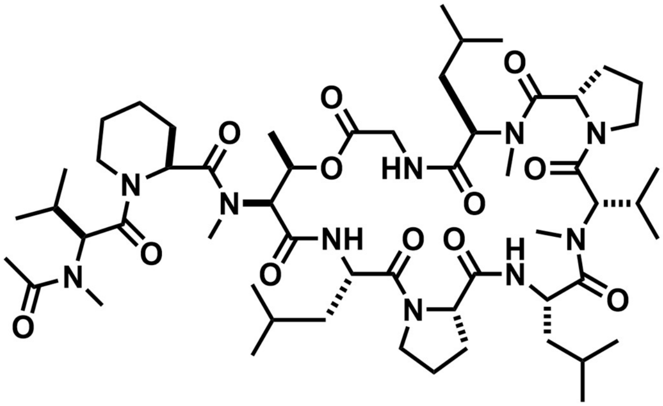

Chemical formula: $C_{56}H_{94}N_{10}O_{12}$. Exact mass (Da): 1098.705. m/z calculated for $[M + Na]^+$: 1121.695; found 1020.861.

MALDI data shown is from the peak collected at 21 minutes. The product does not appear stable under standard RP-HPLC conditions and was not assayed for antibacterial activity.

**24**

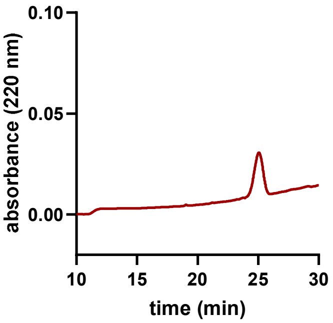


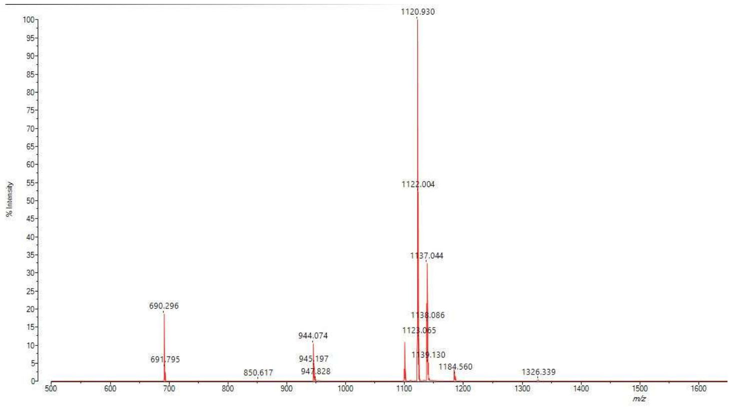


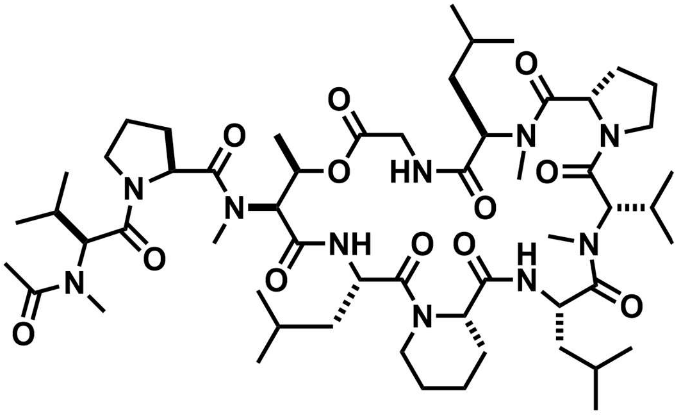

Chemical formula: $C_{56}H_{94}N_{10}O_{12}$. Exact mass (Da): 1098.705. m/z calculated for $[M + Na]^+$: 1121.695; found: 1120.903.

**25**

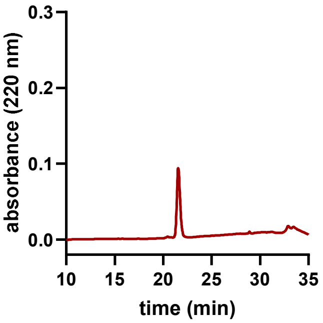


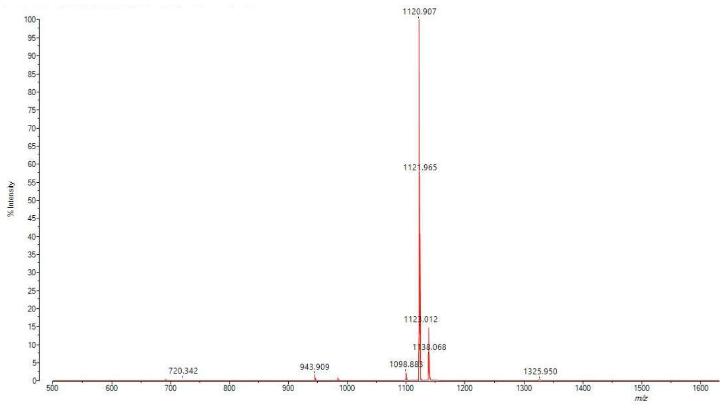

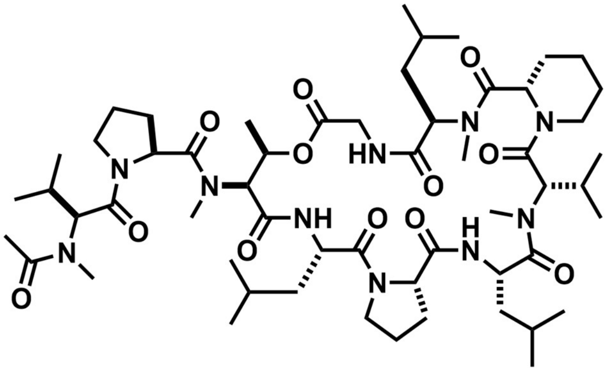

Chemical formula: $C_{56}H_{94}N_{10}O_{12}$. Exact mass (Da): 1098.705. m/z calculated for $[M + Na]^+$: 1121.695; found: 1120.907.

**26**

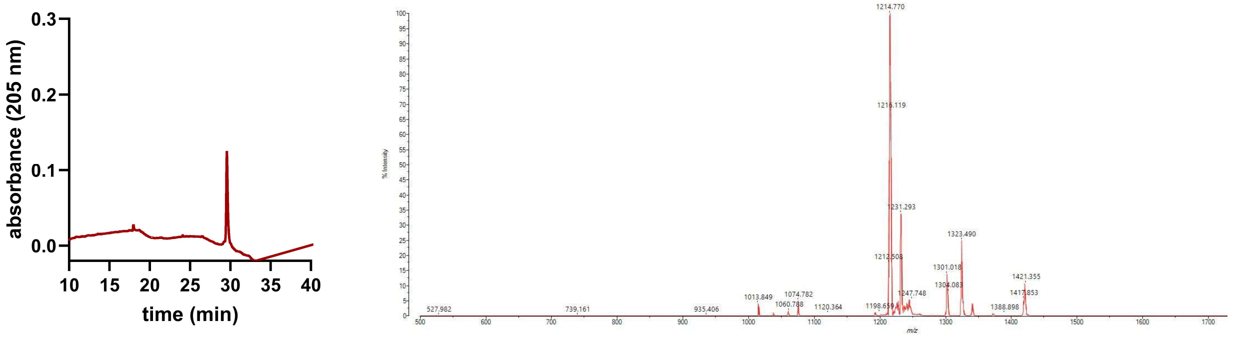


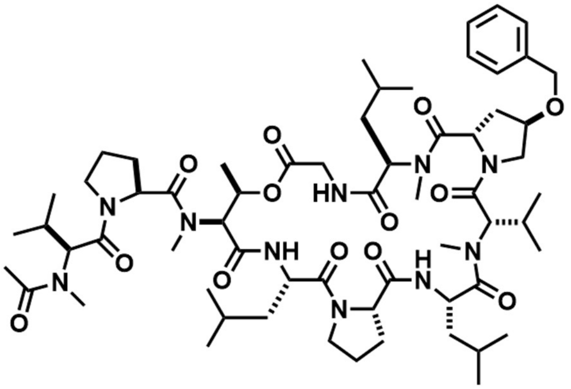

Chemical formula: $C_{62}H_{98}N_{10}O_{13}$. Exact mass (Da): 1190.731. m/z calculated for $[M + Na]^+$: 1213.721; found: 1214.770.

**27**

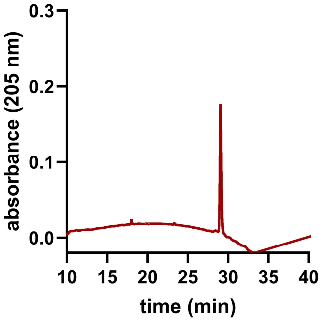


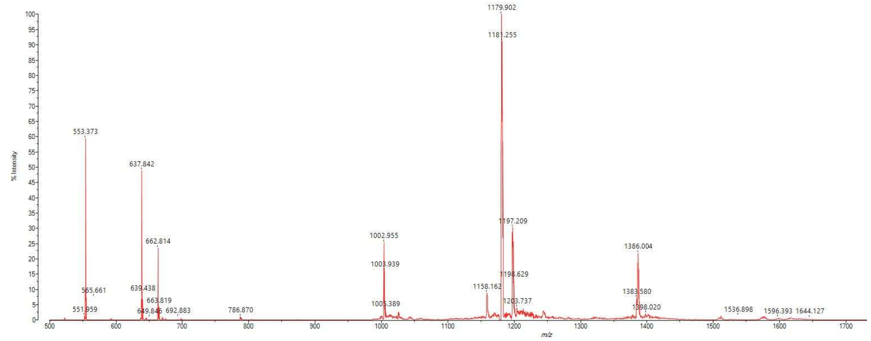

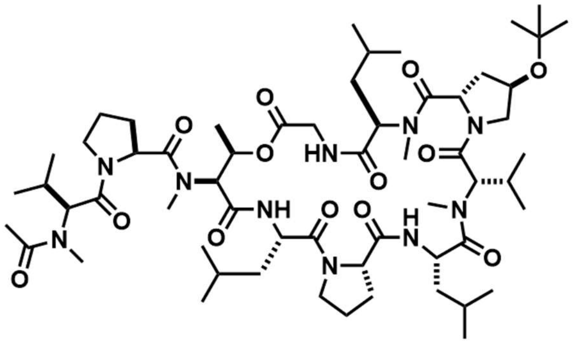

Chemical formula: $C_{59}H_{100}N_{10}O_{13}$. Exact mass (Da): 1156.747. m/z calculated for $[M + Na]^+$: 1179.737; found: 1179.902.

**28**

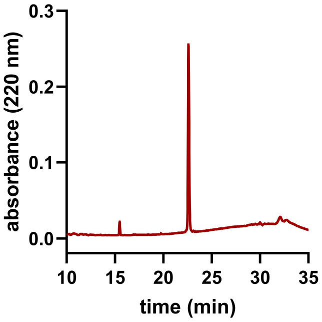


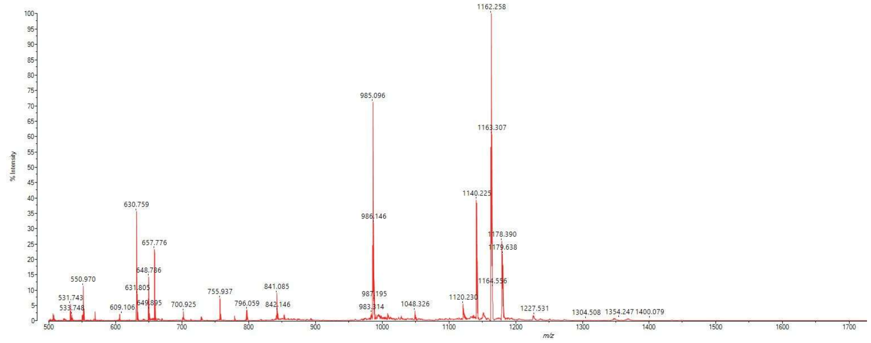

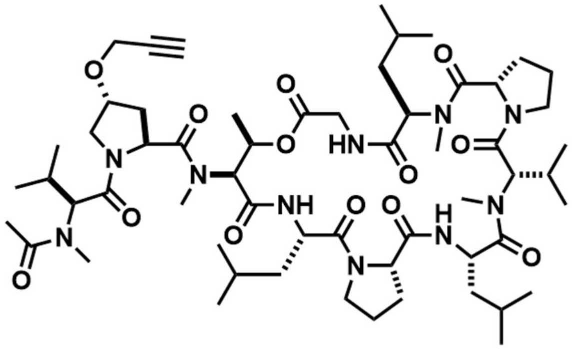

Chemical formula: $C_{59}H_{94}N_{10}O_{13}$. Exact mass (Da): 1138.700. m/z calculated for $[M + Na]^+$: 1161.690; found: 1062.258.

**29**

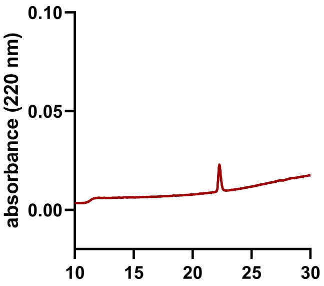


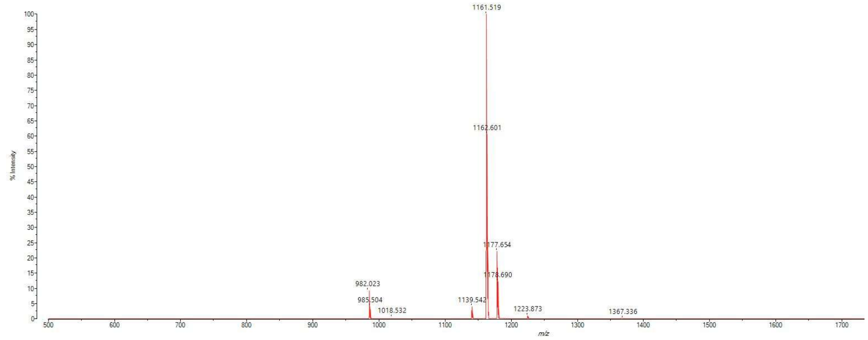


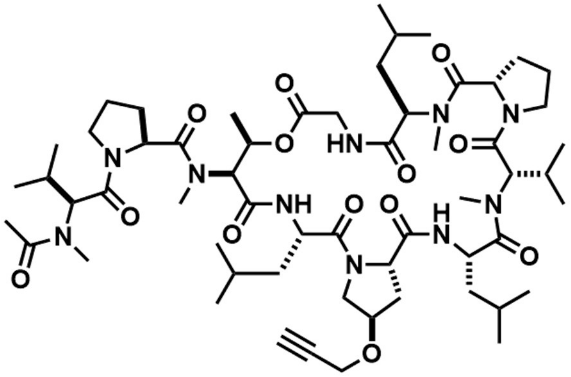


Chemical formula: $C_{59}H_{94}N_{10}O_{13}$. Exact mass (Da): 1138.700. m/z calculated for $[M + Na]^+$: 1161.690; found: 1061.519.

**30**

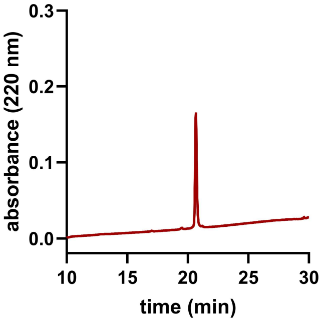


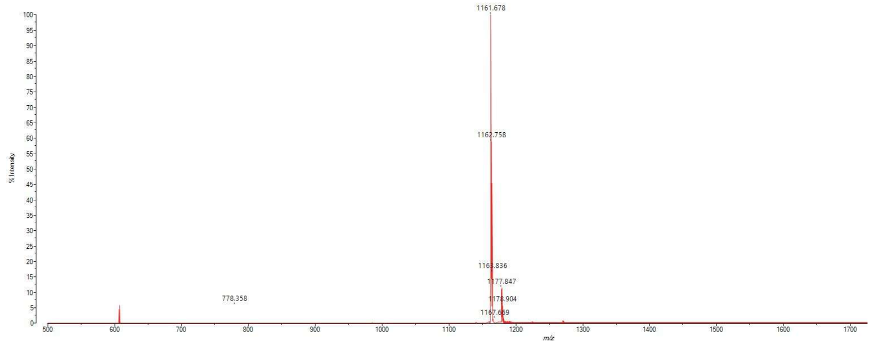


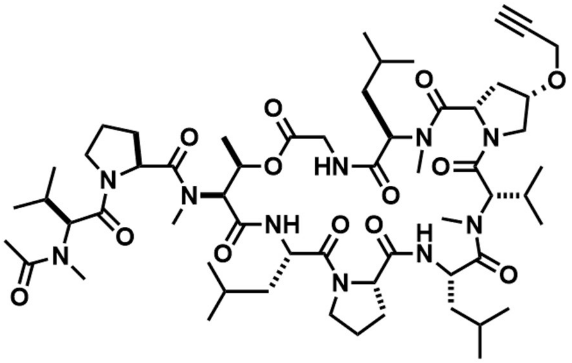


Chemical formula: $C_{59}H_{94}N_{10}O_{13}$. Exact mass (Da): 1138.700. m/z calculated for $[M + Na]^+$: 1161.690; found: 1061.678.

**31**

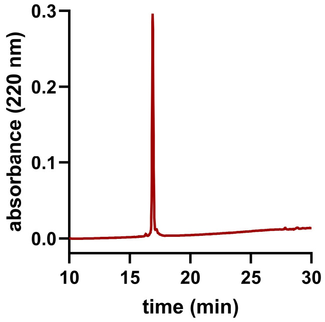


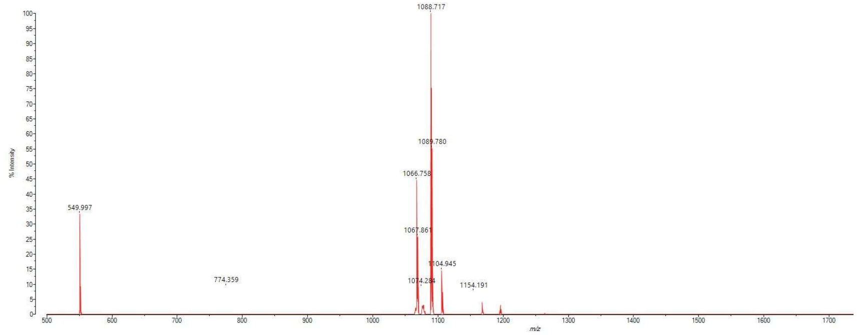


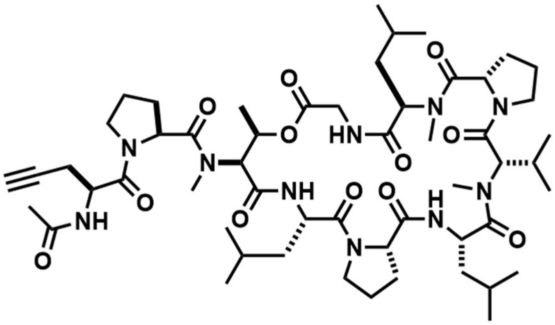

Chemical formula: $C_{54}H_{86}N_{10}O_{12}$. Exact mass (Da): 1066.643. m/z calculated for $[M + Na]^+$: 1089.633; found: 1088.717.

**32**

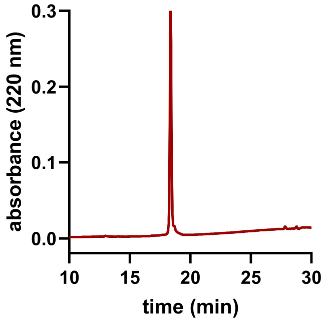


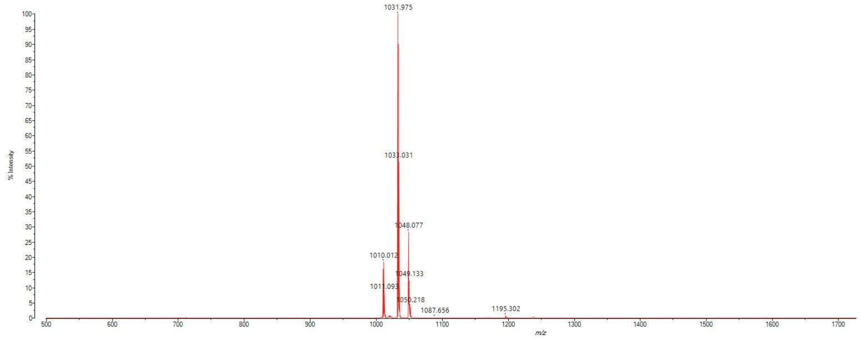


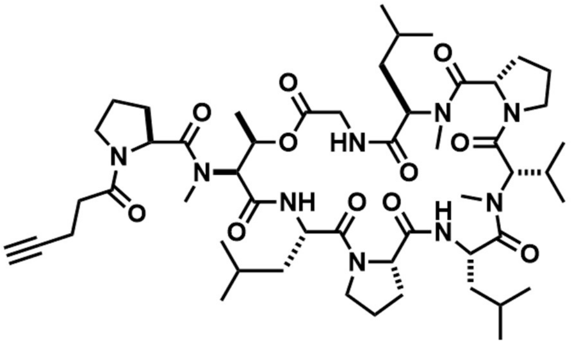

Chemical formula: $C_{52}H_{83}N_{9}O_{11}$. Exact mass (Da): 1009.621. m/z calculated for $[M + Na]^+$: 1032.611; found: 1031.975.

**33**

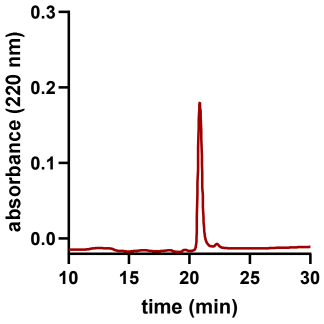


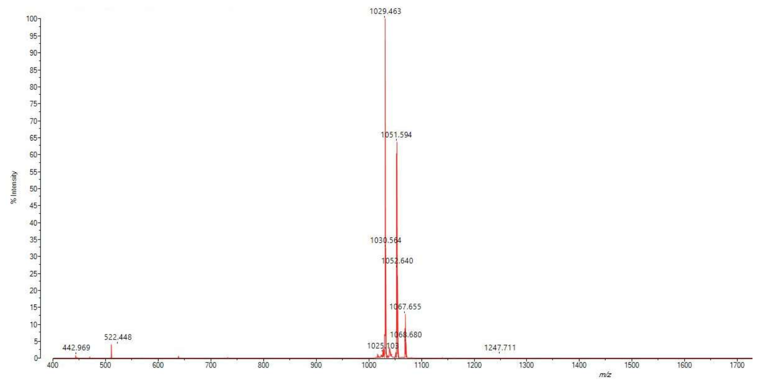


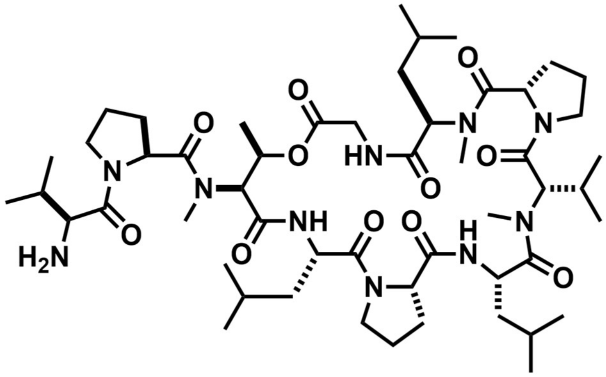

Chemical formula: $C_{52}H_{88}N_{10}O_{11}$. Exact mass (Da): 1028.663. m/z calculated for $[M + H]^+$: 1029.670; found: 1029.463.

**34**

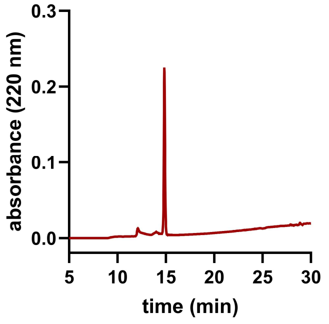


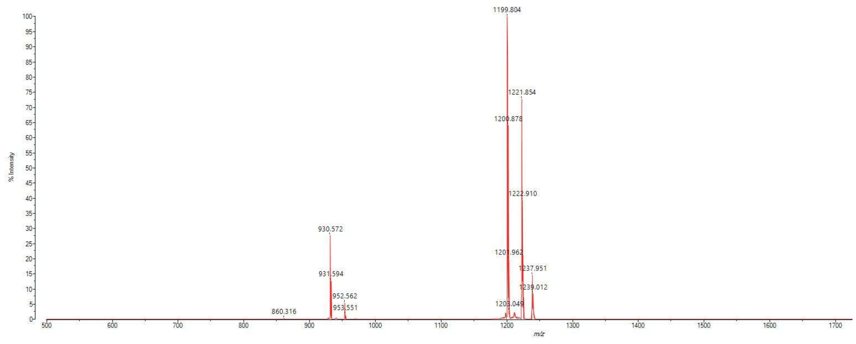

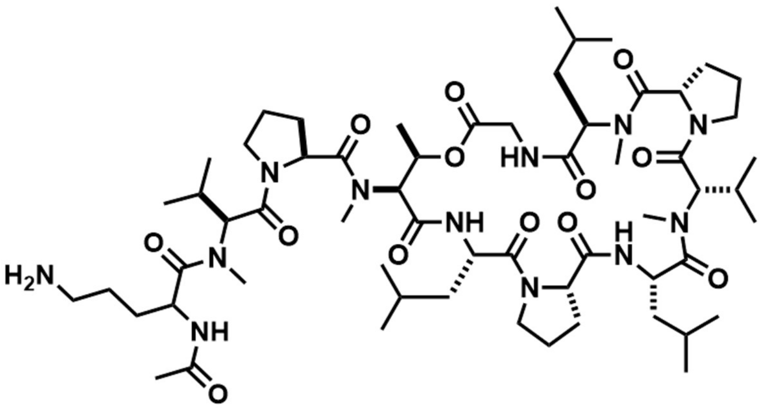

Chemical formula: $C_{60}H_{102}N_{12}O_{13}$. Exact mass (Da): 1198.769. m/z calculated for $[M + H]^+$: 1199.776; found: 1199.804.

**35**

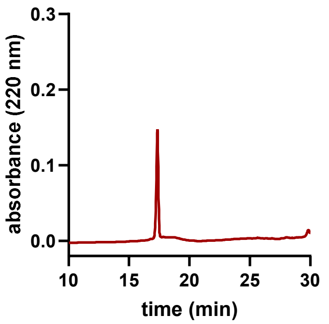


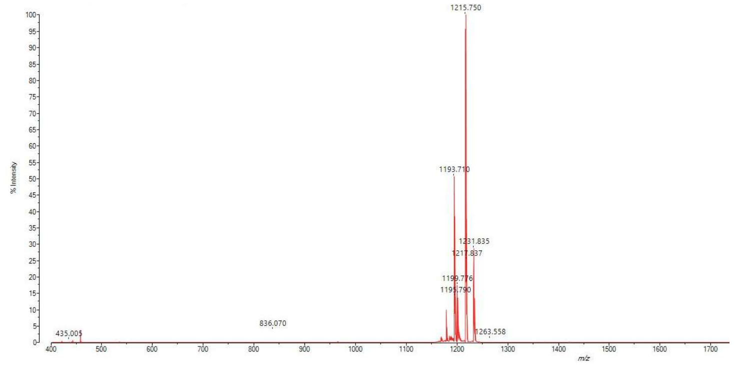


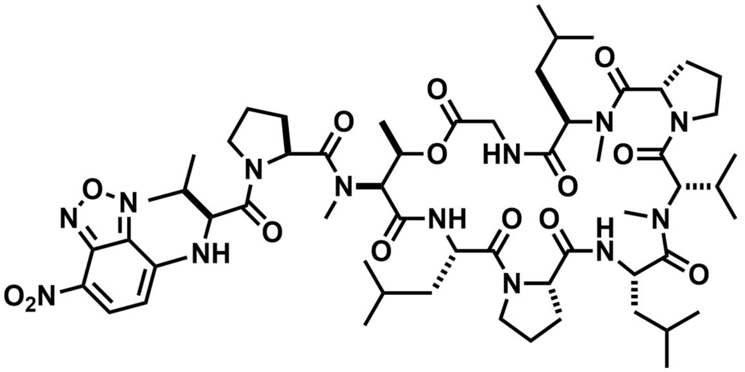

Chemical formula: $C_{58}H_{89}N_{13}O_{14}$. Exact mass (Da): 1191.665. m/z calculated for $[M + H]^+$: 1192.672; found: 1193.710.

**36**

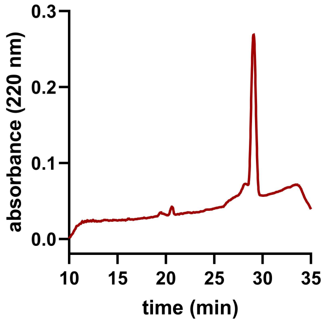


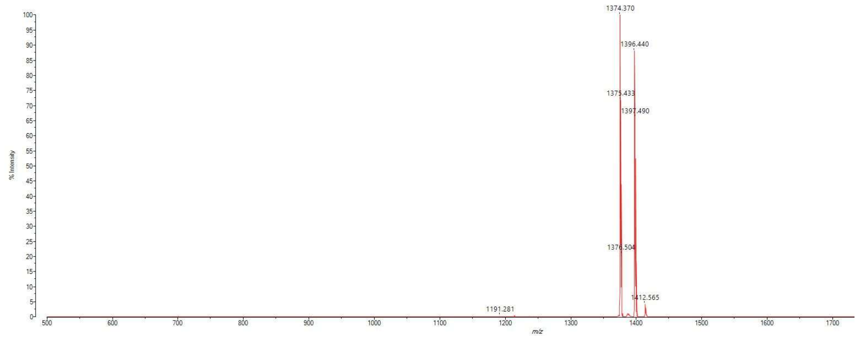


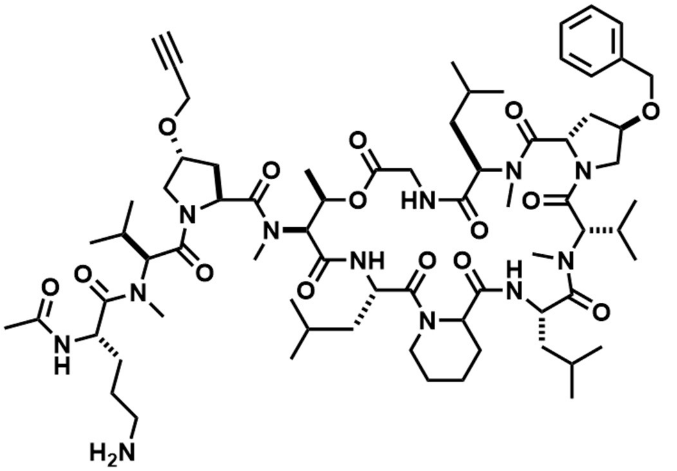


Chemical formula: $C_{71}H_{112}N_{12}O_{15}$. Exact mass (Da): 1372.837. m/z calculated for $[M + Na]^{+}$: 1395.827; found: 1396.440.

### Characterization of *trans*-*N*-Boc-4-propargyloxy-L-proline

$^{1}H$ NMR ($CDCl_3$) 400 MHz

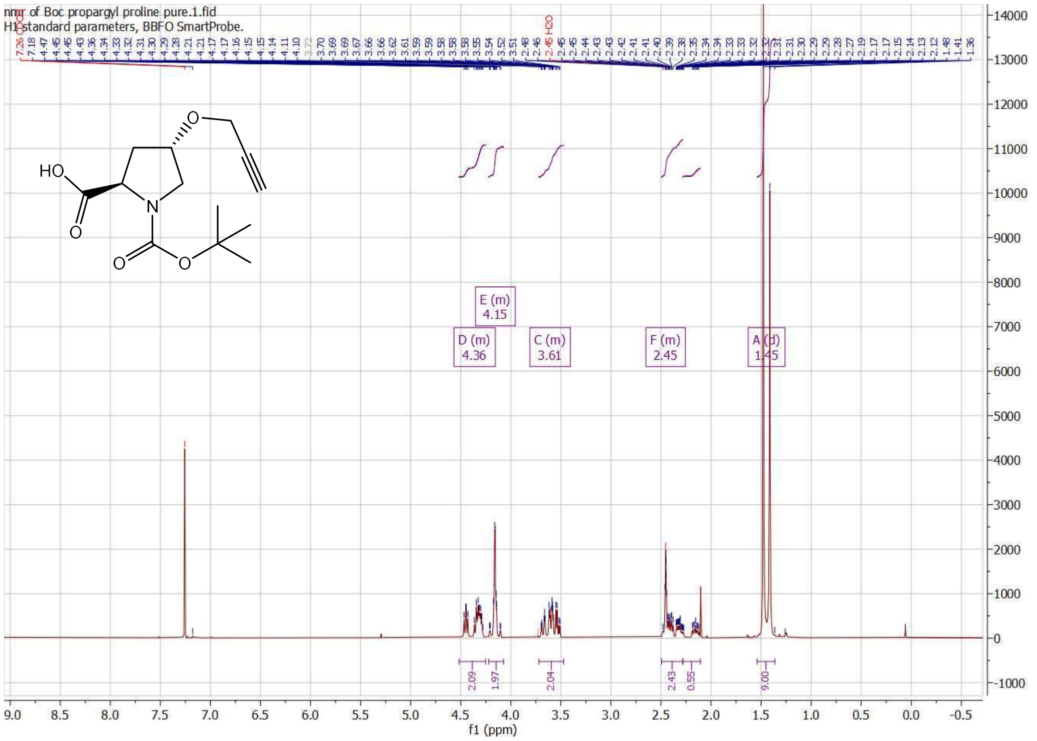


$^{13}C$ NMR ($CDCl_3$) 400 MHz

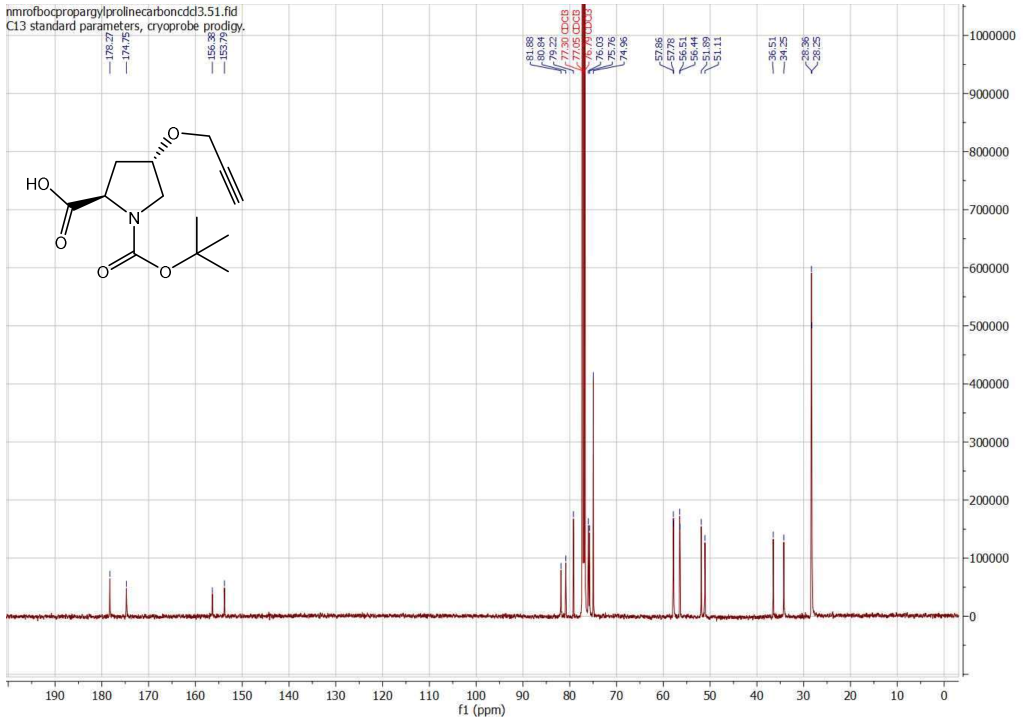

ESI-MS: calculated: 269.30, observed: 269.27 [M-H]$^{-}$

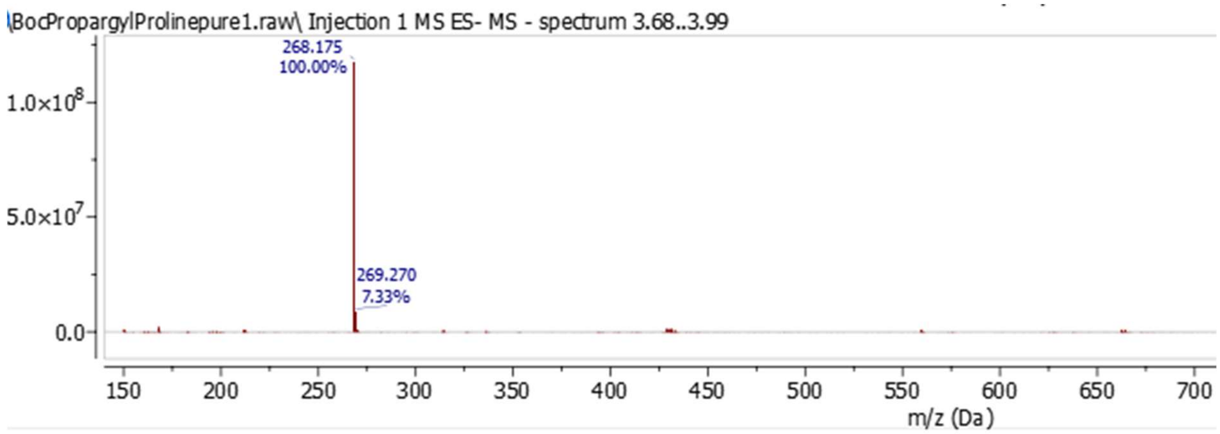

### Characterization of *trans-N*-Fmoc-4-propargyloxy-L-proline

$^{1}$H NMR ($CDCl_3$) 400 MHz

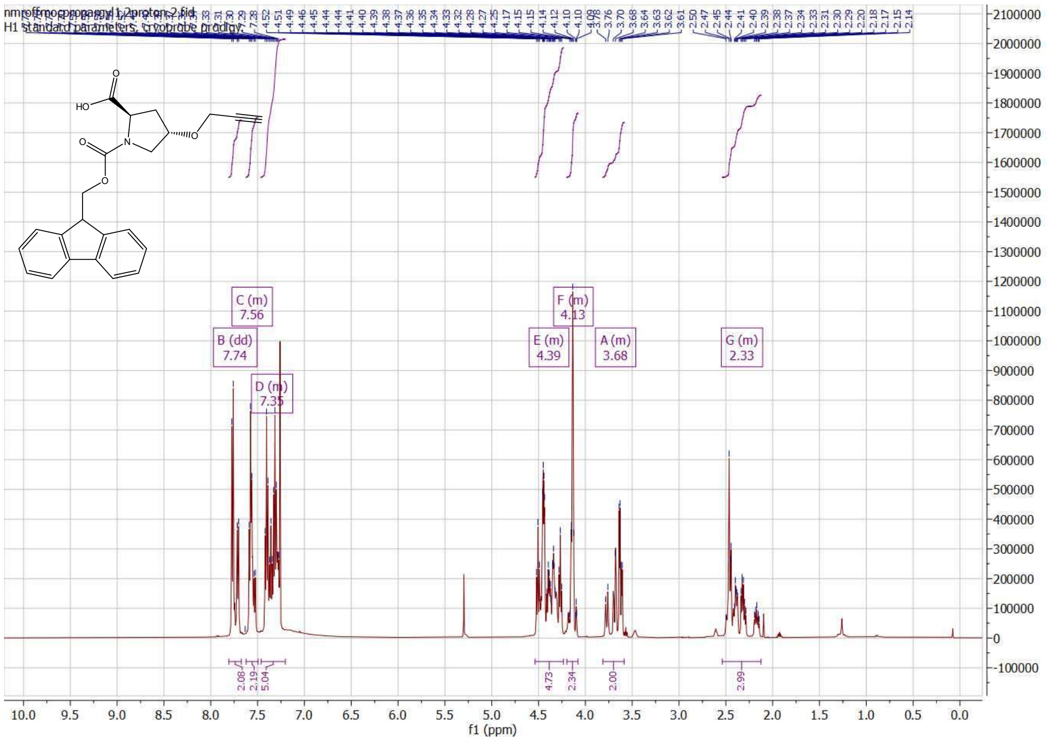


$^{13}$CNMR ($CDCl_3$) 400 MHz

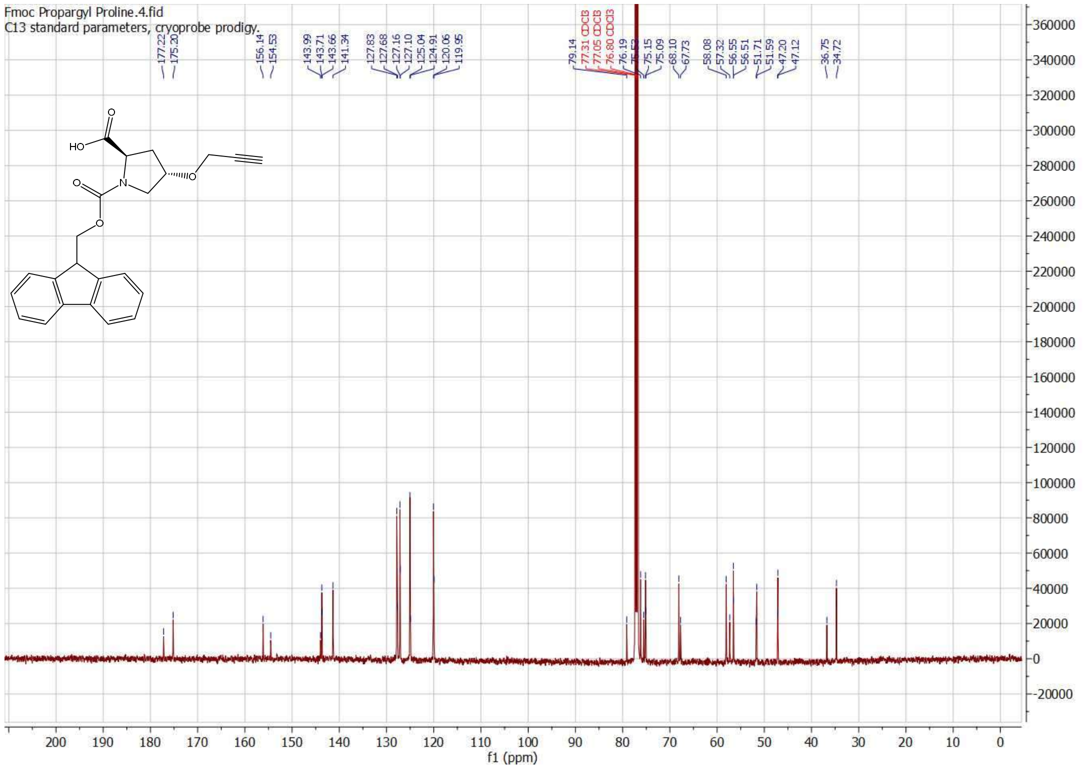

ESI-MS: calculated: 391.42, observed: 414.39 $[M+Na]^+$

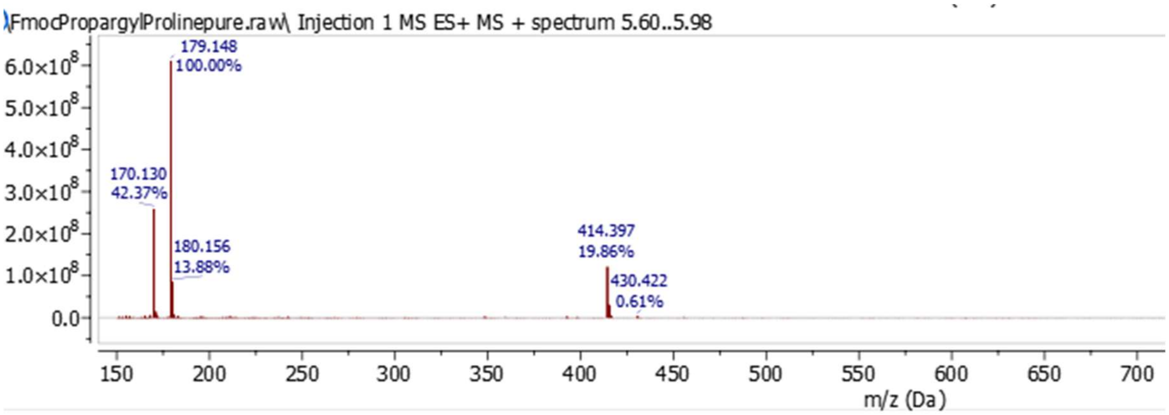


UPLC trace (254 nm):

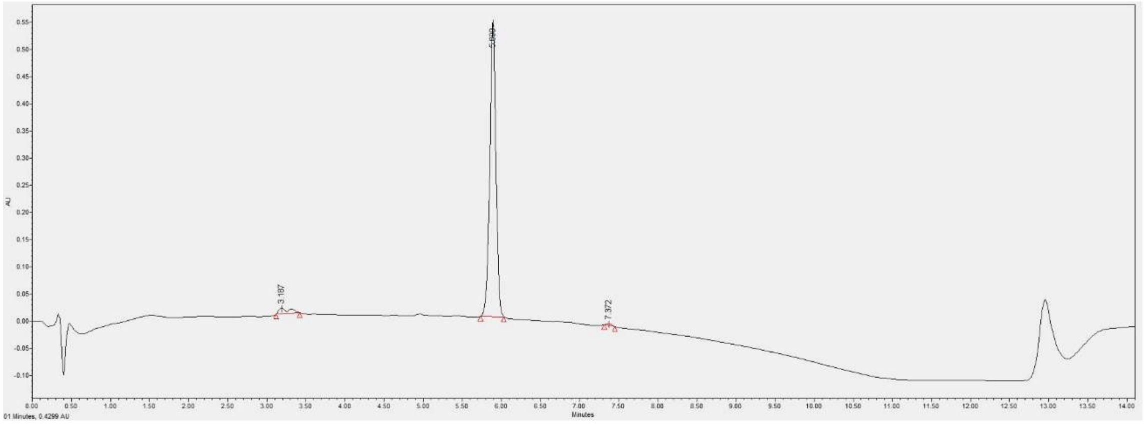
